# Optimized CRISPR-mediated gene knock-in reveals FOXP3-independent control of human Treg identity

**DOI:** 10.1101/2021.01.16.426937

**Authors:** Avery J. Lam, David T.S. Lin, Jana K. Gillies, Prakruti Uday, Anne M. Pesenacker, Michael S. Kobor, Megan K. Levings

## Abstract

Treg cell therapy is a promising curative approach for a variety of immune-mediated conditions. CRISPR-based genome editing allows precise insertion of transgenes through homology-directed repair, but use in human Tregs has been limited. We report an optimized protocol for CRISPR-mediated gene knock-in in human Tregs with high-yield expansion. To establish a benchmark of human Treg dysfunction, we targeted the master transcription factor FOXP3 in naive and memory Tregs. Although FOXP3-knockout Tregs upregulated cytokine expression, effects on suppressive capacity manifested slowly and primarily in memory Tregs. Moreover, FOXP3-knockout Tregs retained their characteristic phenotype and had few changes in their DNA methylation landscape, with FOXP3 maintaining methylation at regions enriched for AP-1 binding sites. Thus, while FOXP3 is important for human Treg development, it has a limited role in maintaining mature Treg identity. Optimized gene knock-in with human Tregs will enable mechanistic studies and the development of tailored, next-generation Treg cell therapies.

## Introduction

Regulatory T cells (Tregs) are critical mediators of immune tolerance and homeostasis. Decades of research into their use as a cell therapy have highlighted their preclinical potential as a curative treatment for a variety of immune-mediated conditions, reflected in the multitude of completed and ongoing clinical trials of Treg transfer in autoimmunity, graft-versus-host disease, and solid organ transplantation (Ferreira et al., 2019). While these trials have found Treg cell therapy to be safe and feasible, their clinical efficacy remains inconclusive in part due to small study sizes (Ferreira et al., 2019). Furthermore, whether these cells will transition into pathological effector T cells in vivo during heightened inflammation as they have in mouse models (Zhou et al., 2009; Bailey-Bucktrout et al., 2013; Komatsu et al., 2014) and human autoimmune disease (Dominguez-Villar, Baecher-Allan and Hafler, 2011; McClymont et al., 2011) is unclear. To address these concerns, a range of work is focused on uncovering fundamental mechanisms controlling Treg identity, fitness, and behaviour, and creating ways to endow therapeutic Tregs with stable functional profiles.

The advent of precision genome editing with CRISPR/Cas9 is a major advance towards realizing the use of targeted gene tailoring to study human Treg biology and create optimal cell therapy products. Early efforts to deliver CRISPR/Cas9 into primary human T cells by plasmids (Mandal et al., 2014; Hendel et al., 2015; Su et al., 2016), lentiviruses, or adenoviruses (Wang et al., 2014; Li et al., 2015; Chen et al., 2018; Shifrut et al., 2018) were hampered by toxicity, low efficiency, and the risk of off-target Cas9 activity (Zhao et al., 2006; Hornung and Latz, 2010). More recently, transfection with pre-formed CRISPR/Cas9 ribonucleoproteins (RNP) by electroporation has emerged as a more efficient approach, with editing rates of 20–90% (Hendel et al., 2015; Schumann et al., 2015; Seki and Rutz, 2018) that also enables multiplexed targeting (Liu et al., 2017; Ren et al., 2017).

Moreover, in the presence of a DNA template with homology to the Cas9 cut site, homology-directed repair (HDR) can mediate seamless gene modification or targeted transgene integration. CRISPR-mediated HDR has been used in primary human T cells to correct point mutations or replace the endogenous TCR with an engineered TCR or chimeric antigen receptor (Eyquem et al., 2017; Roth et al., 2018). In terms of methods, HDR templates have been delivered to human T cells via ssDNA, dsDNA, or adeno-associated virus (AAV), with the latter achieving consistently higher targeting efficiencies (40–70%) (Sather et al., 2015; Gwiazda et al., 2016; Eyquem et al., 2017) than with naked DNA (10–50%) (Roth et al., 2018; Nguyen et al., 2020), due in part to the HDR-promoting properties of AAV (Hirata and Russell, 2000; Vasileva, Linden and Jessberger, 2006).

The use of CRISPR and CRISPR-mediated HDR in primary human Tregs has so far been limited. CRISPR/Cas9 editing in human Tregs with either RNP or lentivirus yielded a range of editing efficiencies (30–80%) dependent on target and gRNA (Chen et al., 2018; Noel et al., 2018; Seki and Rutz, 2018; Schumann et al., 2020). Combining CRISPR with a HDR ssDNA template, one group reported partial restoration of suppressive capacity in Tregs by correcting an IL2RA mutation (Roth et al., 2018). More recently, another group used AAV to insert a FOXP3 cDNA in CD4^+^ T cells as a therapeutic strategy to override deleterious FOXP3 mutations. The resulting HDR-edited cells could be enriched with a reporter gene and exhibited partially restored suppressive function, though HDR rates in Tregs were low (10–20%) (Goodwin et al., 2020).

While these results are promising, human Tregs have a comparatively lower expansion potential than CD4^+^CD25^−^ conventional T cells (Tconv), and, depending on the purity of the starting population, can be prone to spontaneous destabilization and death upon extended in vitro culture (Hoffmann et al., 2006, 2009; Marek et al., 2011; Schmidl et al., 2011; Hansmann et al., 2012). Finding ways to maximize the yield of genome-edited human Tregs to enable in-depth functional studies and therapeutic applications requires systematic optimization of CRISPR-mediated HDR efficiency in parallel with maximal in vitro Treg expansion. Here, focusing on editing efficiency and Treg yield, we report an optimized method for gene knock-in in human Tregs to generate a uniform population of gene-edited cells. As a proof of concept, we targeted the master transcription *FOXP3* because of its critical role in mouse and human Treg development and function (Ono, 2020), enabling us to establish a benchmark of human Treg dysfunction at the DNA methylation, phenotypic, and functional levels. This optimized CRISPR-based gene knock-in system for human Tregs will herald not only mechanistic studies but also the development of edited, next-generation Treg cell therapies.

## Results

### Optimization of CRISPR and HDR delivery to human Tregs

Since HDR typically occurs during cell division (Liu et al., 2018) and previous reports of CRISPR editing in T cells involved TCR-mediated pre-activation, we first compared the efficiency of five different CD4^+^ T cell activation methods: (1) Dynabeads Human T-Expander; (2) Dynabeads Human Treg Expander; (3) plate-bound anti-CD3 with soluble anti-CD28 (pCD3 + sCD28); (4) ImmunoCult Human CD3/CD28 T Cell Activator (CD3/CD28 Tetramer), and (5) ImmunoCult Human CD3/CD28/CD2 T Cell Activator (CD3/CD28/CD2 Tetramer). Both Dynabead variants induced strong, early activation, as judged by upregulation of the T cell activation markers CD69, CD25, CD71, and CTLA-4 within 24–72 h, with negligible differences between T-Expander and Treg Expander versions. In contrast, upregulation of these proteins by CD3/CD28 Tetramers or pCD3 + sCD28 was slower and, at the time points examined, lower than with either Dynabead variant (**Figure S1A**). At day 5, sustained expression of LAP, a selective marker of activated Tregs (Tran et al., 2009), was consistently highest with CD3/CD28/CD2 Tetramers, an increase not seen with CD3/CD28 Tetramer (**Figure S1A**). CD4^+^ T cells began to proliferate (Ki-67^+^) 72 h after activation, with a consequent increase in fold expansion on day 5 (**Figure S1A-B**). Since differences in cell yield between the activation reagents were negligible, we elected to pre-activate cells for 5 days to increase cell numbers and selected three activation reagents for further testing in human Tregs.

To optimize CRISPR editing using Tregs, we compared editing efficiency in cells pre-activated with Dynabeads T-Expander, pCD3 + sCD28, or CD3/CD28/CD2 Tetramer using two electroporation voltages. For convenient assessment of protein knockout (KO) rates by flow cytometry, we used a gRNA targeting *CD226* (**Figure 1A**). Due to the limiting numbers of Tregs that can be obtained from blood, we found that Dynabead removal prior to electroporation resulted in substantial cell loss and insufficient cell recovery after editing (data not shown); this was not a limitation with Tconvs (**Figure S1C-D**). Focusing on the remaining conditions, we found that CD3/CD28/CD2 Tetramer pre-activation paired with 1400 V / 30 ms / 1 pulse yielded the optimal balance between editing efficiency and cell recovery, especially compared to a higher electroporation voltage (2000 V / 20 ms / 2 pulses) (**Figure 1B**). For Tconvs, both Dynabeads and CD3/CD28/CD2 Tetramer yielded high editing efficiency and cell recovery (**Figure S1C-D**). Differences in cell recovery were due to electroporation and not CD226 KO (**Figure S1E**).

**Figure 1.**
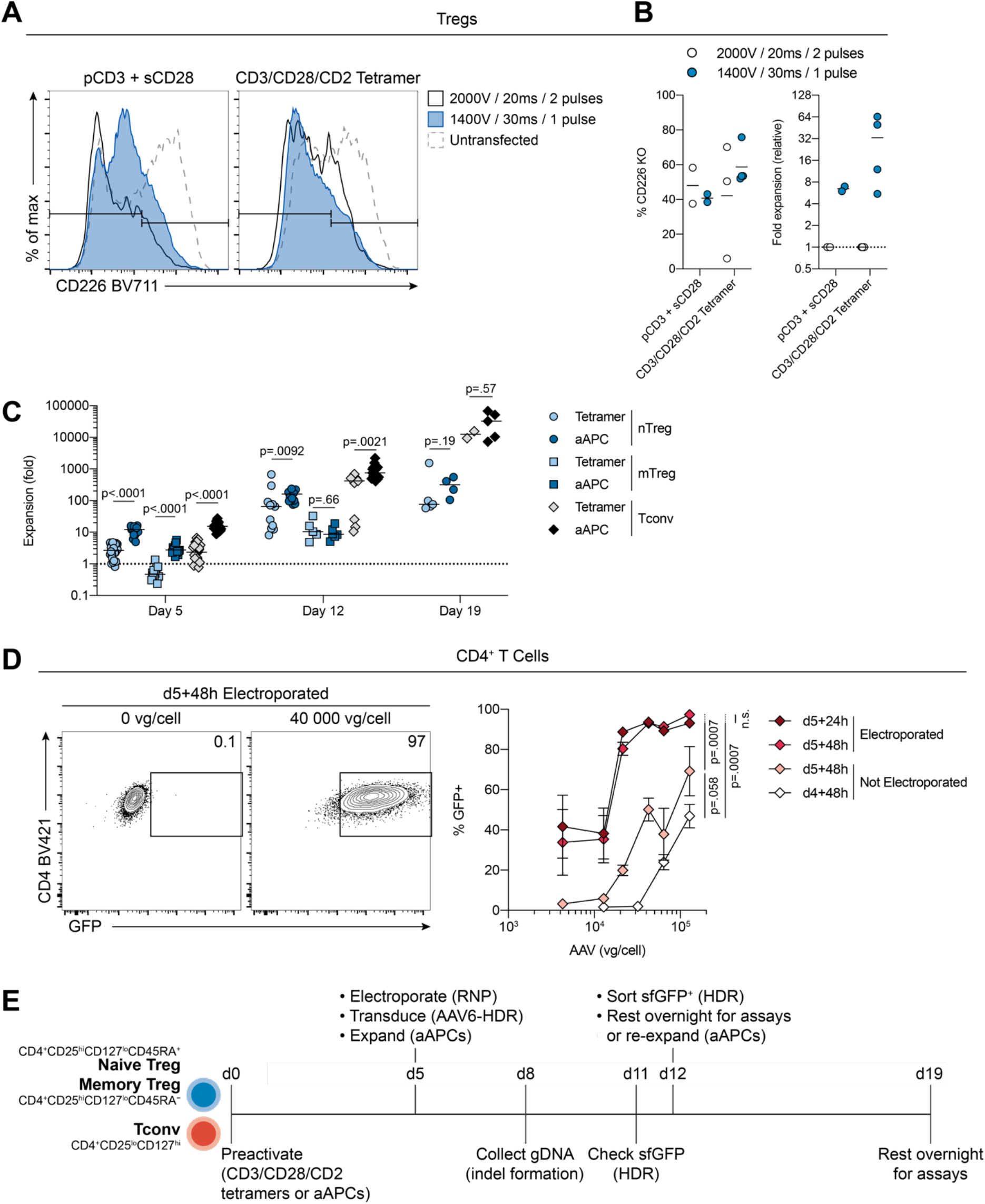
Optimization of CRISPR and AAV delivery to human Tregs. (**A-B**) Tregs (CD4^+^CD25^hi^CD127^lo^) were pre-activated for 5 d with the indicated reagent, electroporated with a *CD226*-targeting gRNA, then expanded for a further 3 d with artificial antigen-presenting cells (aAPCs). (**A**) Representative CD226 expression. (**B**) Left: percent CD226 KO, relative to untransfected cells. Right: fold expansion, relative to 2200 V / 20 ms / 2 pulses within each activation agent. (n=2–4, 2 experiments). (**C**) Naive Tregs (nTreg; CD4^+^CD25^hi^CD127^lo^CD45RA^+^), memory Tregs (mTreg; CD4^+^CD25^hi^CD127^lo^CD45RA^−^), and conventional T cells (Tconv; CD4^+^CD25^lo^CD127^hi^) were pre-activated with the indicated method (5 d), electroporated with Cas9, then expanded by restimulation with aAPCs every 7 d. Fold expansion over time (n=2–24, 1–22 experiments). Due to low expansion, mTregs were not quantified at d 19. (**D**) CD4^+^ T cells were pre-activated with CD3/CD28/CD2 tetramers (4–5 d), electroporated or not, transduced with AAV6-CMV-GFP, and expanded with aAPCs (24–48 h) (or not). Representative (left) and quantified (right) GFP expression. (**E**) Schematic diagram of optimized human Treg expansion with CRISPR/HDR-mediated gene knock-in. (**B**) Solid lines and dots represent means and individual donors, respectively; (**C**) depicts median. Significance determined by (**C**) Mann-Whitney test (nTregs, mTregs, and Tconvs evaluated separately) and (**D**) 1-way ANOVA with Tukey’s multiple comparisons test of the areas under the curve. See also Figure S1.

Tregs represent only a small proportion of peripheral blood CD4^+^ T cells (<5%), so substantial in vitro expansion is needed for in vitro study and therapeutic application. Moreover, naive and memory Tregs (nTregs and mTregs, defined by the presence or absence of CD45RA, respectively) have distinct characteristics (Miyara et al., 2009) and expansion potential (Hoffmann et al., 2006, 2009; Marek et al., 2011). We thus investigated the ability of CD3/CD28/CD2 tetramers to expand nTregs vs. mTregs in comparison to an artificial antigen-presenting cell (aAPC)-based method we previously found to be optimal for nTreg expansion (Dijke et al., 2016; MacDonald et al., 2019). aAPC-based pre-activation resulted in a substantial and consistent increase in nTreg and mTreg yield at day 5 (**Figure 1C**). This advantage was maintained at day 12 for nTregs (median (interquartile range): 162-fold (98–201) with aAPCs vs. 65-fold (13–91) with tetramers); all pre-activation conditions were expanded on aAPCs from day 5 onwards. In contrast, mTregs, having limited expansion potential (Hoffmann et al., 2006, 2009; Marek et al., 2011), exhibited little difference in yield on day 12 and were not further quantified at day 19 (**Figure 1C**).

Finally, to optimize HDR delivery, we used AAV6 (Sather et al., 2015; Wang et al., 2016) encoding a constitutively expressed GFP. A wide range of AAV6 titres and delivery time points have been reported for HDR editing in human T cells, from 10^3^ to 10^6^ viral genomes (vg)/cell and addition within minutes to hours after electroporation (Sather et al., 2015; Gwiazda et al., 2016; Wang et al., 2016; Eyquem et al., 2017; Hale et al., 2017). We compared two approaches: (1) pre-transducing with AAV6 one day before electroporation (i.e., day 4), such that cells would have the AAV6-HDR template available at the time of electroporation on day 5, or (2) transducing with AAV6 immediately after electroporation (day 5). In CD4^+^ T cells, saturation of transduction occurred within 24 h and was maintained for at least 48 h (**Figure 1D**). Transducing cells on day 4 yielded low GFP expression, with a moderate improvement if AAV6 was delivered on day 5 without electroporation. In contrast, electroporation significantly increased AAV6 transduction (**Figure 1D**), likely due to bypassing the requirement for AAV receptor-mediated entry.

Overall, critical parameters for maximizing the yield of gene-edited Tregs included T cell pre-activation reagent and duration, the choice of electroporation voltage, and timing and mode of AAV transduction. **Figure 1E** depicts our optimized expansion timeline for Treg subsets with CRISPR-mediated HDR editing. Briefly, flow-sorted Tregs are pre-activated with CD3/CD28/CD2 tetramers or aAPCs for 5 days, then electroporated (1400 V / 30 ms / 1 pulse) with a CRISPR/Cas9 RNP, immediately transduced with AAV6-HDR, and restimulated with aAPCs for 7 days. At day 12, HDR-edited Tregs are purified based on reporter gene expression, then either rested overnight in reduced IL-2 for functional assays or restimulated with aAPCs for further expansion.

### Efficient CRISPR/HDR-mediated gene knock-in in human Treg subsets

Having established an optimized protocol to edit Tregs with CRISPR and HDR, we functionally validated our system with *FOXP3* as a proof-of-concept gene. We designed five gRNAs targeting relatively 5′ *FOXP3* exons not subject to alternate splicing (**Figure S2A**) and tested their editing efficiency in nTregs. Genomic editing was evaluated 3 days after electroporation since the DNA repair process takes 48–72 h to complete (Brinkman et al., 2018). FOXP3 CR1, CR3, and CR5 gRNAs achieved high protein ablation rates (**Figure 2A-B**). *FOXP3* is located on the X chromosome and is subject to random inactivation in female somatic cells (Tommasini et al., 2002; Carrel and Willard, 2005; Cotton et al., 2011): as such, genomic indel formation rates upon CRISPR editing in Tregs from females were approximately half of those from males (**Figure 2C**). Nevertheless, FOXP3 protein loss was comparable regardless of sex (**Figure 2A-B**). CR1 and CR3 targeted 5′ regions of *FOXP3* (**Figure S2A**) and were hence suitable for developing gene knock-in strategies.

**Figure 2.**
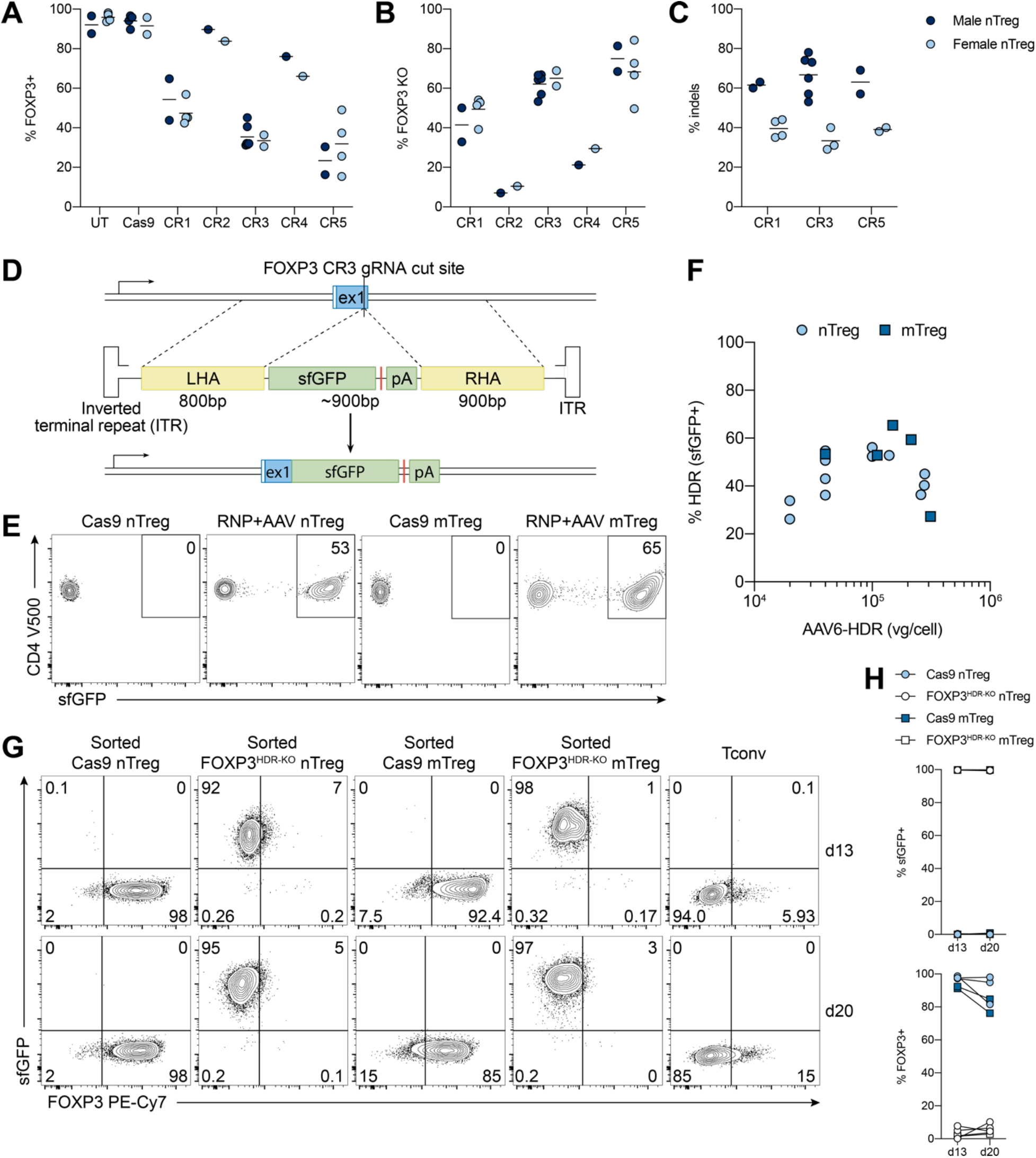
Efficient CRISPR/HDR-mediated gene knock-in in human Treg subsets. (**A-C**) nTregs were pre-activated with CD3/CD28/CD2 tetramers (5 d), edited with the indicated *FOXP3*-targeting gRNA, and expanded with aAPCs (3 d) (n=6 male, n=5–6 female, 4–7 experiments). (**A**) FOXP3 expression, by flow cytometry. (**B**) Percent FOXP3 KO, relative to Cas9 or untransfected (UT) cells. (**C**) Percent indel formation. (**D**) Schematic diagram of promoterless FOXP3 HDR design and targeted integration. Red line (stop codon). (**E-F**) nTregs and mTregs were pre-activated with CD3/CD28/CD2 tetramers (5 d), electroporated with FOXP3 CR3 gRNA, transduced with AAV6-HDR, and expanded with aAPCs (6 d) (n=5–8, 3–4 experiments). (**E**) Representative sfGFP (HDR) expression from cells edited and transduced with AAV6-HDR (14×10^4^ viral genomes (vg)/cell). (**F**) sfGFP (HDR) in nTregs and mTregs. (**G-H**) 7 d after editing, cells were flow-sorted as sfGFP^+^, expanded with aAPCs (7 d), then rested overnight. (**G**) Representative and (**H**) quantified sfGFP and FOXP3 expression (n=3 nTreg, n=2 mTreg, 1–2 experiments). Dots in (**A-C**) and (**H**) represent individual donors. Dots in (**F**) represent individual donor-AAV6 titre conditions. See also Figure S2.

To achieve CRISPR/HDR-mediated gene knock-in, we included an in-frame reporter transgene (superfolder GFP; sfGFP) flanked by two homology arms (∼800 bp each) in the HDR template design, such that targeted integration results in truncation of *FOXP3* exon 1 and *sfGFP* knock-in (**Figure 2D**). Because AAV stably persists in cells as episomes, this promoterless HDR design ensures sfGFP expression faithfully reports HDR editing and avoids the risk of exogenous-promoter driven upregulation of nearby genes (Lombardo et al., 2011). We achieved 40–50% gene knock-in (sfGFP^+^) in nTregs and 40–60% in mTregs, with an AAV titre of at least 4×10^4^ vg/cell (**Figure 2E-F**). Increases in HDR editing efficiency were not seen beyond AAV titres of 1×10^5^ vg/cell (**Figure 2F**). The addition of a commercially available HDR-enhancing reagent did not substantially increase HDR efficiency (**Figure S2B**). Interestingly, HDR editing increased overall FOXP3 KO rates by 10–25% relative to CRISPR editing alone (**Figure S2C**), suggesting that HDR-mediated gene disruption increases gene-KO efficiency.

We sorted HDR-edited nTregs and mTregs as sfGFP^+^ cells and expanded them for an additional 7 days. Sort-purified, HDR-edited Tregs (hereon FOXP3^HDR-KO^) were uniformly FOXP3-ablated and sfGFP^+^, and this phenotype was stable over time (**Figure 2G-H**). In contrast, Cas9 nTregs and mTregs maintained high FOXP3 expression in this 2–3-week timeframe (**Figure 2G-H**). Because of the promoterless HDR design, sustained sfGFP expression suggested that *FOXP3* transcription remained active in human Tregs even in the absence of FOXP3 protein, consistent with some reports in mice (Gavin et al., 2007; Lin et al., 2007) but not others (Bending et al., 2018).

### High transgene expression distinguishes HDR editing from episomal AAV6

We designed a second HDR template to include an exogenous promoter-driven transgene to accommodate genes not constitutively and/or uniformly expressed in Tregs. Specifically, the design incorporated the EF-1α promoter, which minimally disrupts proximal gene expression upon targeted integration at multiple loci (Lombardo et al., 2011), and a truncated nerve growth factor receptor (ΔNGFR) reporter, a clinically-compatible surface molecule that can be magnetically selected by antibody-bound beads (**Figure 3A**). With this design, AAV6-transduced Tregs could express ΔNGFR without genomic integration. Indeed, when compared to untransduced Tconvs, ΔNGFR expression was maintained in AAV6-transduced Tregs without CRISPR editing (both nTregs and mTregs) for up to a week (**Figure 3B-C**), indicative of episomal AAV6 persistence. ΔNGFR expression was progressively lost as cell divided, with mTregs maintaining higher levels over time due to their lower proliferative capacity (**Figure 1D, Figure 3B-C**). No ΔNGFR expression driven by episomal persistence could be detected by 12 days post-transduction (**Figure 3C**).

**Figure 3.**
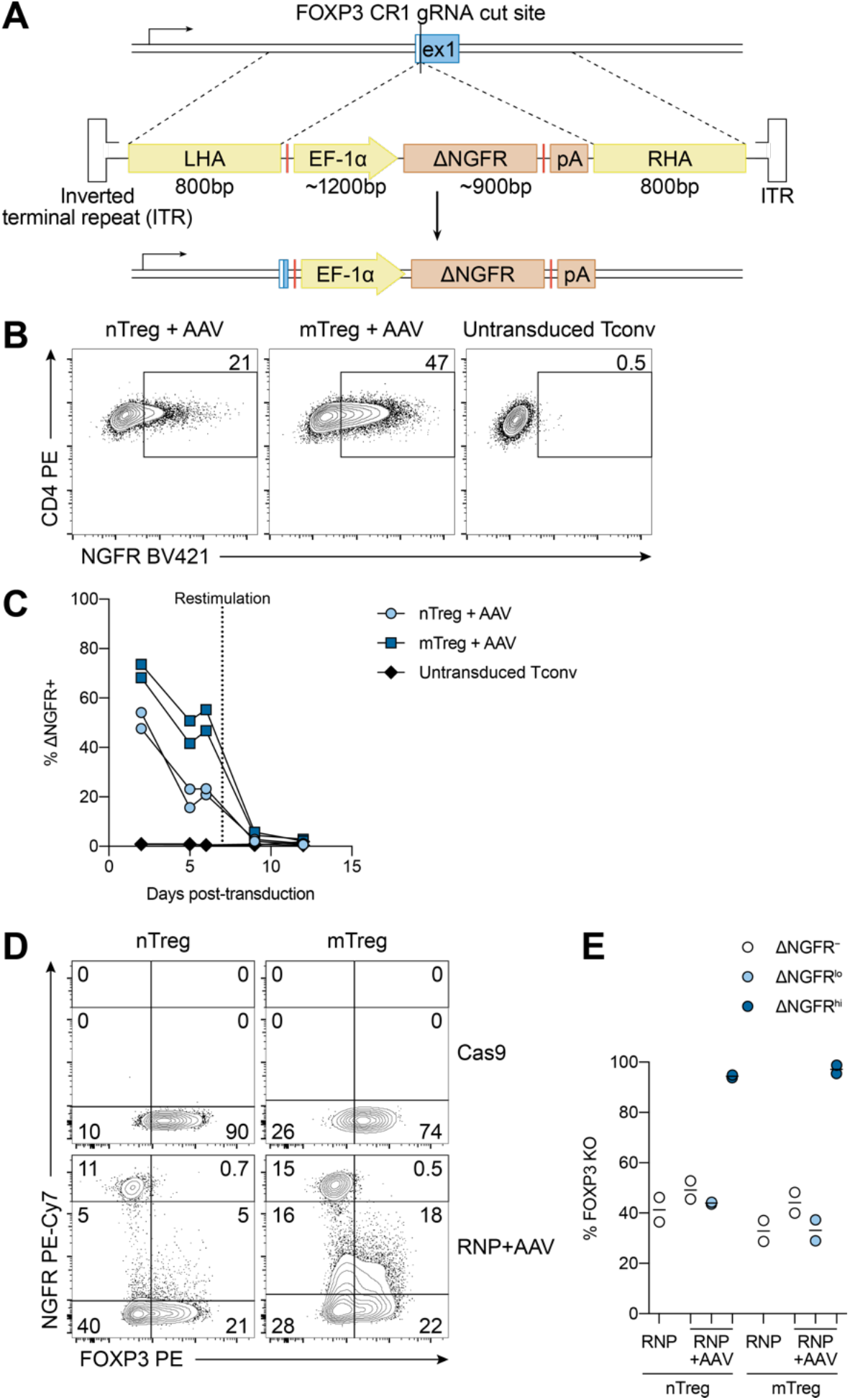
High transgene expression distinguishes HDR editing from episomal AAV6. (**A**) Schematic diagram of promoter-driven FOXP3 HDR design and targeted integration. Red line (stop codon). (**B-C**) nTregs and mTregs were pre-activated with aAPCs (5 d), electroporated without any RNP, transduced with AAV6-HDR (1250 vg/cell), and expanded with aAPCs every 7 d (n=2, 1 experiment). Tconvs expanded in parallel but left untransduced served as ΔNGFR-negative controls. (**B**) Representative ΔNGFR expression 6 d post-transduction. (**C**) ΔNGFR expression over time. (**D-E**) aAPC-pre-activated nTregs and mTregs (5 d) were electroporated with FOXP3 CR1 gRNA, transduced with AAV6-HDR (3600 vg/cell), and expanded with aAPCs (6 d) (n=2, 1 experiment). (**D**) Representative ΔNGFR and FOXP3 expression 6 d post-editing. (**E**) Cells were gated as ΔNGFR^−^, ΔNGFR^lo^, and ΔNGFR^hi^. Percent FOXP3 KO, relative to Cas9 cells. Dots in (**C**) and (**E**) represent individual donors; solid lines in (**E**) represent means.

Notably, when we combined FOXP3 CRISPR and HDR editing with a promoter-driven ΔNGFR HDR template, HDR-edited cells could be distinguished by their high expression of ΔNGFR (**Figure 3D-E**). ΔNGFR^hi^ nTregs and mTregs exhibited near-complete FOXP3 ablation, while ΔNGFR^lo^ cells showed no such enrichment in FOXP3 KO, relative to cells only edited with FOXP3 gRNA (**Figure 3D-E**). Thus, even with their relatively limited proliferative potential, Tregs can be edited with CRISPR and AAV-based HDR editing containing a constitutive promoter and identified by strong transgene expression.

### FOXP3^HDR-KO^ promotes Treg secretion of multiple cytokines but not of IL-2

Having generated uniformly FOXP3-ablated human Tregs, we aimed to establish a benchmark of human Treg dysfunction. We focused on sort-purified FOXP3^HDR-KO^ Tregs generated with a promoterless HDR template to limit the risk of transcriptional deregulation at *FOXP3*-proximal elements if inserting a constitutive promoter. We hypothesized that FOXP3 ablation in mature human Tregs would elevate cytokine production because it operates in cooperation or competition with other factors such as NFAT, RUNX1, and RORγt to repress inflammatory cytokine transcription, including *Ifng, Il2, Il4*, and *Il17a* (Bettelli, Dastrange and Oukka, 2005; Wu et al., 2006; Ono et al., 2007; Zheng et al., 2007; Ichiyama et al., 2008; Zhou et al., 2008). Following TCR activation for 4 days, both naive and memory subsets of FOXP3^HDR-KO^ Tregs secreted higher levels of Th1, Th2, and Th17 cytokines (**Figure 4A**). Differential cytokine production was not due to differential proliferation, as KO and Cas9 control Tregs divided comparably in this timeframe (**Figure S3**). FOXP3^HDR-KO^ nTregs predominantly upregulated IFN-γ, IL-5, and IL-13; an increase in the Th17-associated cytokine IL-17A was exclusive to FOXP3^HDR-KO^ mTregs (**Figure 4A**). FOXP3-ablated mTregs also secreted more of the pleiotropic cytokines IL-6, IL-9, and IL-10 compared to their naive counterparts (**Figure 4A**). Thus, FOXP3 ablation in mature Tregs promotes the secretion of a variety of cytokines in a Treg subset-dependent manner.

**Figure 4.**
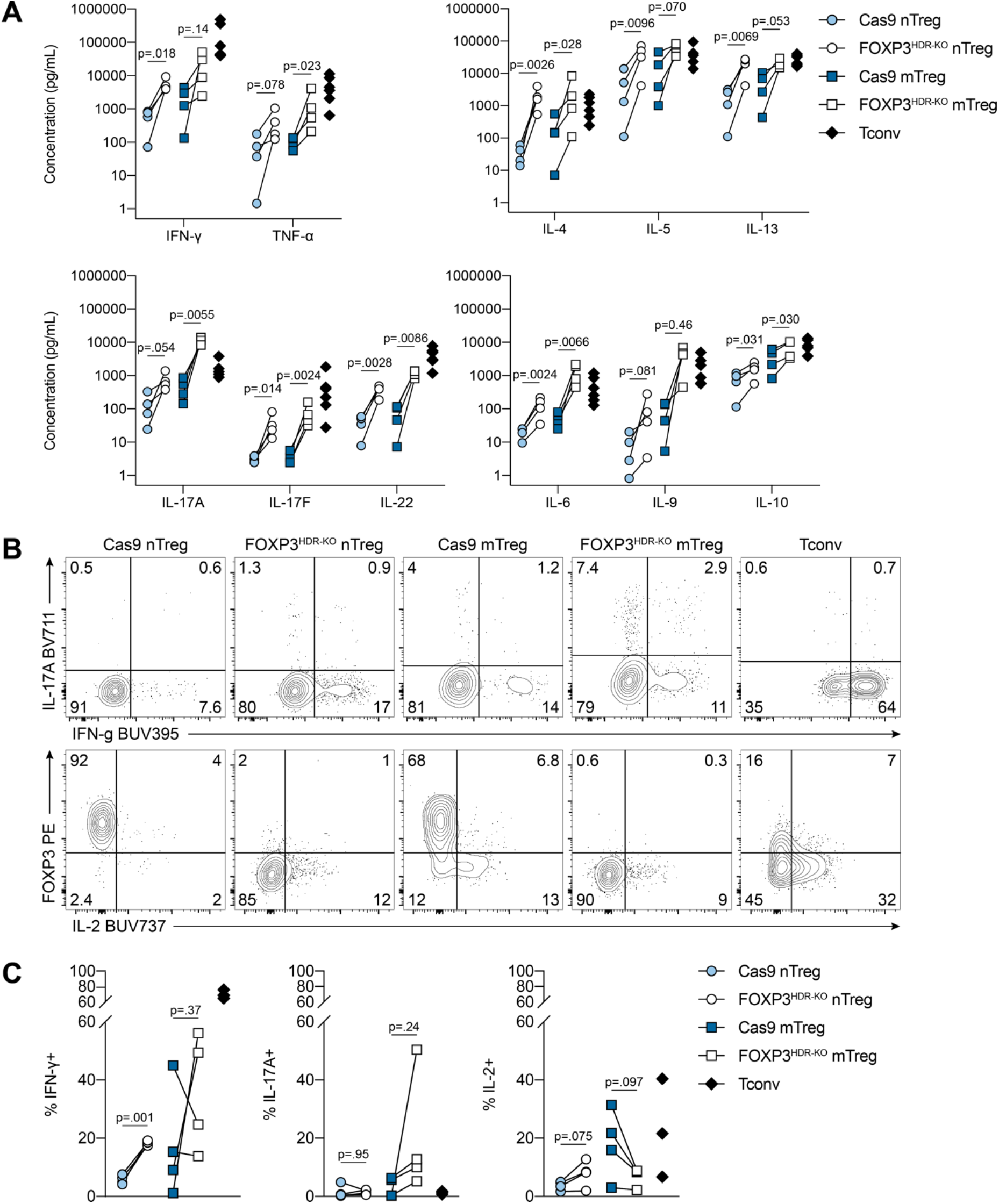
FOXP3^HDR-KO^ promotes secretion of multiple cytokines but not of IL-2. (**A-C**) Cas9 and FOXP3^HDR-KO^ nTregs and mTregs were expanded for 13 d total (n=4, 3 experiments). (**A**) Cytokine concentrations from the supernatants of TCR-activated cells (4 d). (**B-C**) Cells from (**A**) were restimulated with PMA, ionomycin, and brefeldin A (4 h). (**B**) Representative and (**C**) quantified IFN-γ, IL-17A, and IL-2 expression (n=4, 3 experiments). (**A**) and (**C**) depict individual donors. Significance determined by ratio paired t-test in (**A**) and paired t-test in (**C**); nTregs and mTregs evaluated separately. Tconv shown for reference. See also Figure S3.

We also examined cytokine production potential by flow cytometry. Consistent with secreted cytokines in supernatants, FOXP3^HDR-KO^ nTregs upregulated IFN-γ, and there was a trend, though not significant, of IL-17A upregulation by FOXP3^HDR-KO^ mTregs (**Figure 4B-C**). Although lack of IL-2 expression is a hallmark of Tregs and its transcriptional repression is mediated by FOXP3 in cooperation with NFAT (Wu et al., 2006), we did not find elevated IL-2 expression in FOXP3^HDR-KO^ nTreg or mTreg (**Figure 4B-C**), suggesting that additional layers of control beyond FOXP3 suppress IL-2 expression in mature Tregs.

### Subset- and time-dependent effects of FOXP3^HDR-KO^ on Treg suppressive function

We asked whether sort-purified FOXP3^HDR-KO^ Tregs exhibited reduced suppressive capacity in vitro. One week following *FOXP3* editing, there was no change in FOXP3^HDR-KO^ nTreg-mediated suppression of CD4^+^ or CD8^+^ T cells, but FOXP3^HDR-KO^ mTregs were less able to suppress CD8^+^ T cell proliferation, relative to their Cas9 counterparts (**Figure 5A**). Hypothesizing that the downstream effects of transcription factor ablation may manifest progressively over time, we also examined Treg suppressive capacity two weeks after *FOXP3* editing. By this time, FOXP3^HDR-KO^ mTregs were unable to suppress CD4^+^ T cell proliferation, and both FOXP3^HDR-KO^ nTregs and mTregs had reduced abilities to suppress CD8^+^ T cells (**Figure 5B**). Overall, the loss of suppressive capacity of FOXP3-ablated human Tregs was a progressive effect, dependent on Treg subset and responder T cell type: mTregs exhibited a more dramatic loss in suppressive function upon FOXP3 ablation than did nTregs.

**Figure 5.**
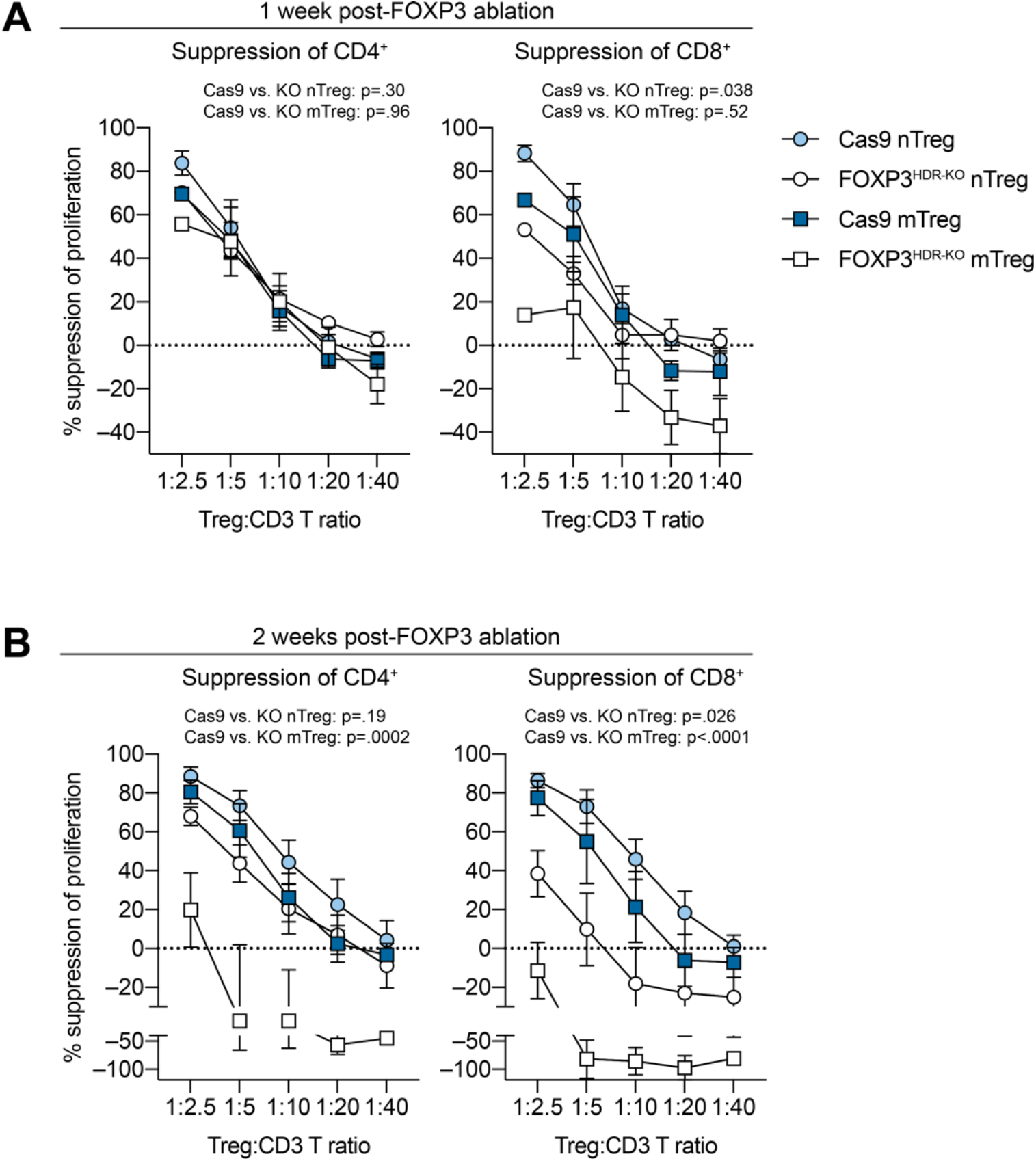
Subset- and time-dependent effects of FOXP3^HDR-KO^ on Treg suppressive function. (**A-B**) Cas9 and FOXP3^HDR-KO^ nTregs and mTregs were expanded for (**A**) 13 d or (**B**) 20 d total (3 experiments). Suppression of proliferation of CD4^+^ (left) and CD8^+^ (right) T cells ((**A**) n=2–4 nTreg, n=1–3 mTreg; (**B**) n=4–5 nTreg, n=3 mTreg). (**A-B**) depict mean±SEM. Significance determined by t-test of the areas under the curve; nTregs and mTregs evaluated separately. See also Figure S4.

Alterations in suppressive function were likely not due to altered Treg viability, as Tregs were detectable at most Treg:CD3^+^ T cell ratios and FOXP3^HDR-KO^ Tregs maintained sfGFP expression, both one week and two weeks after FOXP3 ablation (**Figure S4A-B**). In fact, the less suppressive FOXP3^HDR-KO^ Tregs were more proliferative than their Cas9 counterparts in the assay (**Figure S4A-B**).

We also examined Treg-mediated suppression of cytokines by CD3^+^ T cells, focusing on IFN-γ, TNF-α, and IL-22 because these were secreted by Tregs at substantially lower levels than Tconvs (**Figure 4A**). One week after editing, both nTregs and mTregs were able to suppress cytokine secretion in a dose-dependent fashion, regardless of FOXP3 ablation (**Figure S4C**). In line with their loss of suppression of proliferation, two weeks after editing FOXP3^HDR-KO^ mTregs, but not nTregs, lost their ability to suppress the secretion of IFN-γ and IL-22 (**Figure S4D**). While we cannot exclude the possible contribution of FOXP3^HDR-KO^ mTreg-derived IFN-γ and IL-22, the highest cytokine concentrations were not observed in the condition containing the most mTregs, so it is unlikely that Tregs are the primary source of cytokines in the assay.

### FOXP3^HDR-KO^ Tregs retain a Treg phenotype

Many Treg-associated proteins are encoded by FOXP3 target genes (Sadlon et al., 2010; Birzele et al., 2011; Schmidl et al., 2014) and/or involved in Treg suppressive function (Akkaya and Shevach, 2020). Furthermore, overexpression studies have shown direct Foxp3-mediated transactivation of CD25 and CTLA-4, two proteins highly expressed by Tregs (Chen et al., 2006; Wu et al., 2006). We asked whether Tregs exhibited phenotypic changes upon FOXP3 ablation, hypothesizing that such alterations may underlie defects in suppressive capacity.

We examined protein expression two weeks after editing, since a substantial loss of Treg suppression was observed by this time. Unexpectedly, FOXP3^HDR-KO^ nTregs and mTregs maintained high expression of CD25 and CTLA-4 relative to Tconvs (**Figure 6A**). Loss of FOXP3 also did not affect expression of Helios, CD39, or LAP, proteins implicated in Treg stability and suppressive capacity (**Figure 6A**). CCR4 expression, however, was significantly reduced specifically in FOXP3^HDR-KO^ nTregs, whereas CXCR3 and CCR6 remained unchanged (**Figure 6B**).

**Figure 6.**
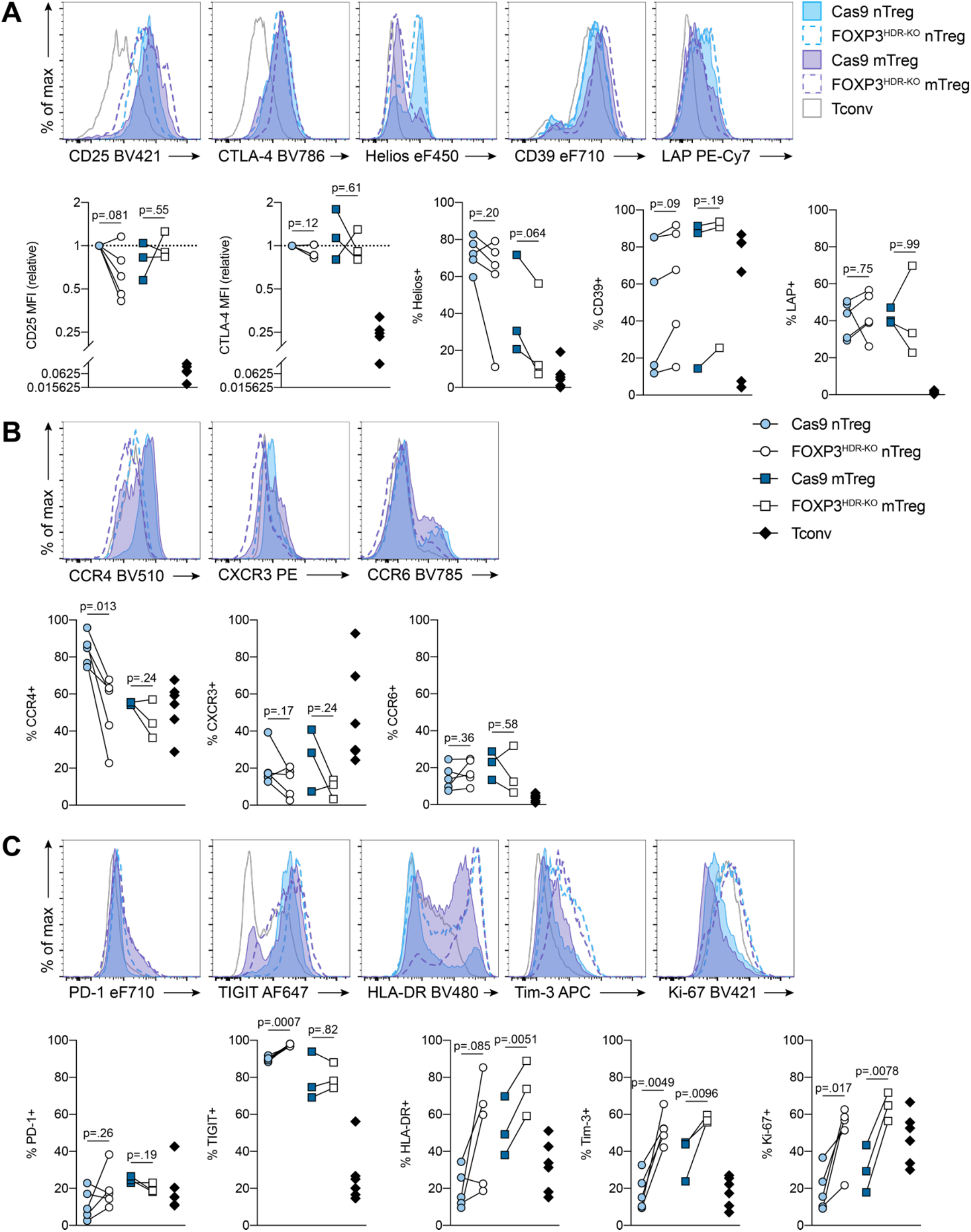
FOXP3^HDR-KO^ Tregs retain a Treg phenotype. (**A-C**) Cas9 and FOXP3^HDR-KO^ nTregs and mTregs were expanded for 20 d total (n=3–5, 3 experiments). (**A**) CD25, CTLA-4, Helios, CD39, LAP, (**B**) CCR4, CXCR3, CCR6, (**C**) PD-1, TIGIT, HLA-DR, Tim-3, Ki-67 expression. (**A-C**) depict individual donors. Significance determined by paired t-test; nTregs and mTregs evaluated separately. Tconv shown for reference. MFI, geometric mean fluorescence intensity.

FOXP3 ablation did not affect PD-1 expression, recently characterized as a Treg-limiting protein (Tan et al., 2021), but did elevate expression of TIGIT (nTregs), HLA-DR (mTregs), and Tim-3 (both nTregs and mTregs) (**Figure 6C**). Although these proteins mark subsets of highly suppressive human Tregs, they are also upregulated upon TCR activation (Baecher-Allan, Wolf and Hafler, 2006; Gautron et al., 2014; Joller et al., 2014; Lucca et al., 2019). FOXP3^HDR-KO^ Tregs were also more proliferative (Ki-67^+^) (**Figure 6C**). These data suggest that FOXP3 ablation exaggerates TCR activation in mature human Tregs.

### FOXP3-independent maintenance of human Treg DNA methylation identity

The overall modest phenotypic changes upon FOXP3 ablation prompted us to investigate its contribution to the maintenance of human Treg identity in lineage-committed cells. A wealth of studies have characterized the distinct molecular signatures of human Tregs at the DNA methylation (Schmidl et al., 2009; Zhang et al., 2013; Ohkura et al., 2020), transcript (Birzele et al., 2011; Ferraro et al., 2014; Pesenacker et al., 2016), and protein levels (Procaccini et al., 2016; Cuadrado et al., 2018). In mice, Foxp3 is necessary but not sufficient to confer Treg identity, especially with respect to DNA methylation (Ohkura et al., 2012; Samstein et al., 2012). Other requisite proteins, such as pioneer transcription factors and enhancers, contribute in tandem (Fu et al., 2012; Rudra et al., 2012; Kitagawa et al., 2017). How the Treg lineage is upheld after Treg development, however, is less well understood (Ono, 2020). To delineate the role of FOXP3 in maintaining human Treg identity after its establishment, we assessed the genome-wide DNA methylation profiles of Cas9 vs. sort-purified FOXP3^HDR-KO^ nTregs as well as Tconvs, all expanded for total 20 days, using the Illumina MethylationEPIC array.

We first performed principal component (PC) analysis across all interrogated CpG sites (702,481) (**Figure 7A**). PC1, explaining 31% of total variance, may be due to inter-individual differences, while PC2, explaining 22% of total variance, clearly segregated nTregs (both Cas9 and FOXP3^HDR-KO^) from Tconvs (**Figure 7A**). We used paired analysis to identify differentially methylated CpG sites between all groups (**Table S1**), finding that FOXP3 ablation induced substantially fewer differentially methylated CpG sites compared to those found between nTregs and Tconvs (**Figure S5A**). Furthermore, unsupervised hierarchical clustering of the most differentially methylated CpG sites revealed that nTregs and Tconvs clustered separately, regardless of FOXP3 ablation (**Figure 7B**). These data suggest that the human Treg DNA methylation landscape is maintained largely independently of FOXP3.

**Figure 7.**
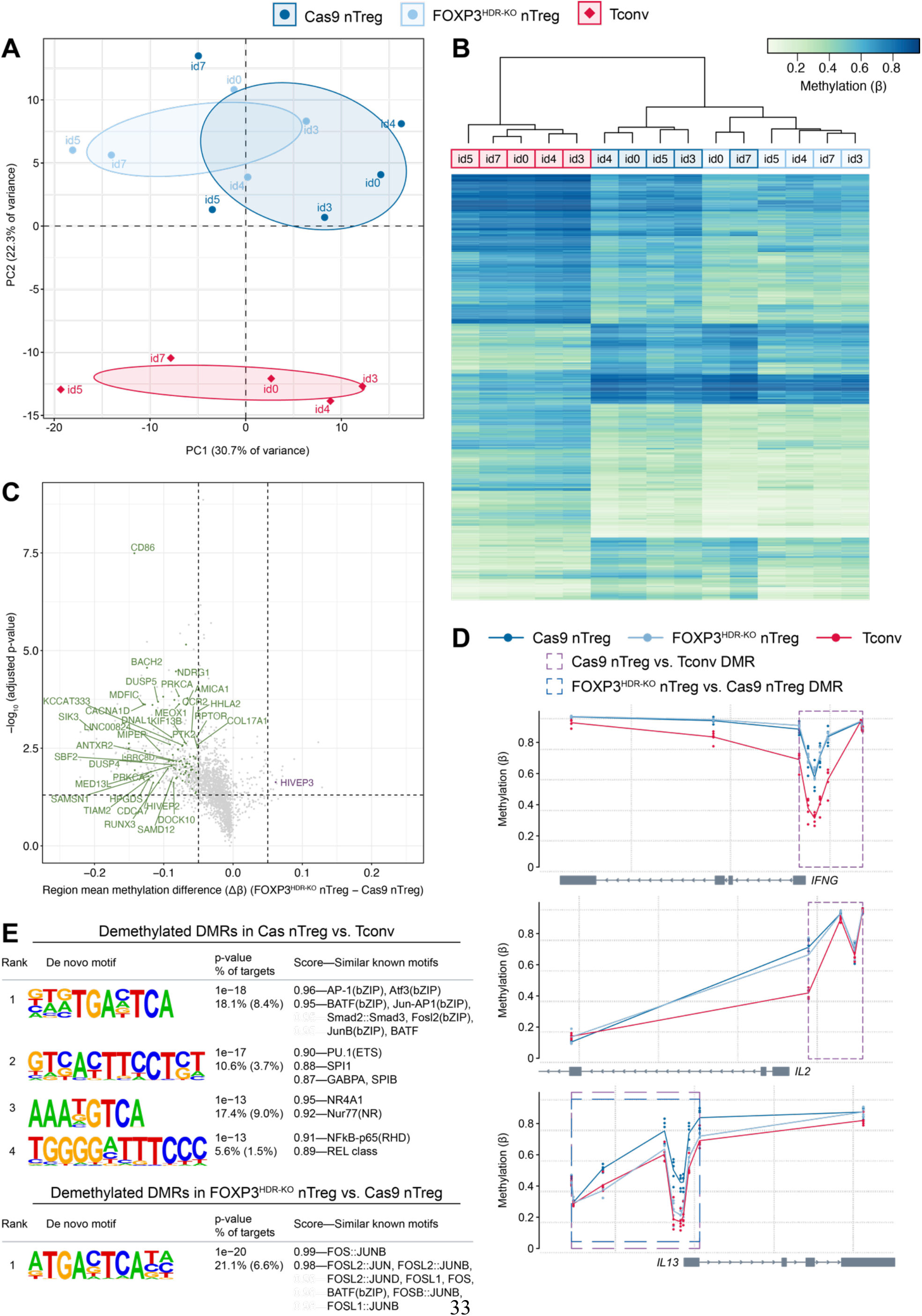
FOXP3-independent maintenance of human Treg DNA methylation identity. (**A-E**) Genome-wide DNA methylation was assessed in 20 d-expanded Cas9 nTregs, FOXP3^HDR-KO^ nTregs, and Tconvs (n=5). (**A**) Principal component analysis across all interrogated CpG sites (702,481). (**B**) Heatmap of the most differentially methylated CpG sites (mean methylation (beta) difference > 0.2, Benjamini-Hochberg-corrected p-value < 0.05). Hierarchical clustering by Ward’s method. (**C**) Volcano plot of DMRs for FOXP3^HDR-KO^ nTregs vs. Cas9 nTregs (region mean methylation (beta) difference > 0.05, Fisher’s multiple-comparison statistic < 0.05). Purple or green colour identifies FOXP3 KO-induced DMRs with higher or lower levels of methylation, respectively, that overlap a FOXP3-binding region; a selection of these DMRs is labelled by their associated gene. (**D**) Gene-associated DMRs for *IFNG, IL2*, and *IL13*. (**E**) Left: de novo motifs enriched (p-value < 1×10^−12^) in Cas9 Treg-specific demethylated DMRs relative to Tconvs (top) and FOXP3 KO-induced demethylated DMRs relative to Cas9 nTregs (bottom), by HOMER. Middle: p-value, % of demethylated DMR sequences containing at least once instance of each motif, and % of background sequences containing the motif (parentheses). Right: similar known motifs, by similarity score. Individual samples are represented as (**A**) dots, (**B**) columns, and (**D**) dots at each CpG site. DMRs are (**C**) represented as dots and delineated by (**C**) dotted lines or (**D**) dotted boxes (purple: Cas9 nTreg vs. Tconv DMR; dark blue: FOXP3^HDR-KO^ nTreg vs. Cas9 nTreg DMR). See also Figure S5 and Table S1.

Concordant methylation changes at adjacent CpG sites are more functionally relevant than individual sites (Vanderkraats et al., 2013; Guo et al., 2017; Schlosberg, VanderKraats and Edwards, 2017; Gatev et al., 2020) so we identified 2,326 Treg-specific differentially methylated regions (DMRs) and 471 FOXP3 KO-induced DMRs (**Table S1**). Compared to Tconvs, Cas9 nTregs exhibited lower methylation of DMRs in Treg signature genes including *CTLA4, IL2RA* (CD25), *IKZF2* (Helios), *IKZF4* (Eos), *LRRC32* (GARP), and *TNFRSF9* (CD137) (**Figure S5B**), consistent with previous reports (Schmidl et al., 2009; Ohkura et al., 2012; Zhang et al., 2013; Nowak et al., 2018). FOXP3 ablation did not significantly alter DNA methylation at these regions, but it did further reduce methylation at the *TIGIT* promoter (**Figure S5B**). These findings are in line with unchanged protein expression of CD25, CTLA-4, and Helios, but upregulation of TIGIT in FOXP3^HDR-KO^ nTregs (**Figure 6A-C**).

Compared to Tconvs, Cas9 nTregs exhibited higher levels of methylation in terms of both individual CpG sites and DMRs (**Figure S5A**). In contrast, FOXP3 ablation predominantly decreased DNA methylation: >95% of FOXP3 KO-induced DMRs were less methylated relative to Cas9 nTregs (**Figure 7C, Figure S5A**). A number of genes associated with these FOXP3 KO-induced demethylated DMRs, including *BACH2, HIVEP2, DUSP4*, and *RPTOR* (mTORC1), have been implicated in mouse Treg development and/or function (**Figure 7C**) (Roychoudhuri et al., 2013; Zeng et al., 2013; Yan et al., 2015; Schumann et al., 2020). We thus used a publicly available FOXP3 ChIP-chip tiled array dataset from human Tregs (Sadlon et al., 2010) to ask if any DMRs overlapped a FOXP3-binding region (**Table S1**). The majority of FOXP3 KO-induced DMRs did not overlap a FOXP3 binding site: only 18% of Cas9 nTreg vs. Tconv and 21% of FOXP3^HDR-KO^ nTreg vs. Cas9 nTreg DMRs overlapped with a FOXP3-binding region. Notably, one FOXP3^HDR-KO^ nTreg-specific DMR with increased methylation relative to Cas9 nTregs was a FOXP3-binding region in *HIVEP3*, a gene previously shown to be differentially methylated in Tregs relative to Tconvs (Zhang et al., 2013) **(Figure 7C**).

Because FOXP3^HDR-KO^ nTregs exhibited elevated cytokine production (**Figure 4A-C**), we investigated whether these changes were reflected at the epigenetic level. Although these cells upregulated IFN-γ (**Figure 4A-C**), FOXP3 ablation did not alter Treg-specific DNA methylation at the *IFNG* promoter (**Figure 7D**). Similarly, DNA methylation at the *IL2* promoter did not differ between FOXP3^HDR-KO^ and Cas9 nTregs (**Figure 7D**), consistent with the lack of effect on IL-2 protein expression (**Figure 4C**). However, in line with elevated IL-5 and IL-13 protein secretion (**Figure 4A**), FOXP3^HDR-KO^ nTregs had decreased promoter DNA methylation at the Th2 cytokines *IL5* and *IL13*, with methylation in these regions resembling Tconvs (**Figure 7D, Figure S5C**). Since neither *IL5* nor *IL13* DMRs overlapped a FOXP3-binding region, altered DNA methylation at Th2 cytokine loci upon FOXP3 ablation operates via an indirect mechanism.

Finally, to better understand the potential functional effects of DMRs on transcription factor activity, we performed transcription factor motif enrichment analysis. In comparison to Tconvs, Cas9 nTreg-specific demethylated DMRs had a statistically significant over-representation of binding sites for bZIP (AP-1), ETS, nuclear receptor, and NF-κB transcription factor families (**Figure 7E**), which are all involved in Treg development and associated with more accessible chromatin regions in mouse Tregs (Isomura et al., 2009; Long et al., 2009; Ruan et al., 2009; Mouly et al., 2010; Polansky et al., 2010; Sekiya et al., 2011, 2013; Samstein et al., 2012; van der Veeken et al., 2020). Interestingly, FOXP3 KO-induced DMRs with decreased methylation were also significantly enriched in motifs for bZIP (AP-1), but not of ETS, nuclear receptor, or NF-κB transcription factor families. These data suggest that FOXP3 contributes to maintenance of DNA methylation at AP-1 motif-enriched regions in a way that is distinct from other Treg-specific demethylated DMRs. As AP-1 preferentially binds unmethylated DNA (Yin et al., 2017), FOXP3 KO-induced aberrant demethylation of these regions may enable AP-1 overactivation.

## Discussion

Here, we used optimized CRISPR/HDR-mediated gene knock-in to generate a homogenous population of FOXP3-ablated human Tregs and study its role in fully lineage-committed naive and memory cells. We found that human FOXP3^HDR-KO^ Tregs retained a Treg phenotype, likely related to FOXP3-independent maintenance of a Treg-characteristic pattern of DNA methylation. Moreover, loss of suppressive function in FOXP3^HDR-KO^ Tregs took time and was primarily evident in mTregs. Overall, these data suggest that FOXP3 may have a limited functional role once Tregs are lineage-committed, particularly in the naive subset. The ability to utilize CRISPR/HDR-mediated gene knock-in in humans Tregs will enable mechanistic interrogation of other genes involved in Treg biology and the targeted integration of therapeutic transgenes for next-generation Treg cell therapies.

Since Tregs are rare and defined by hypo-proliferation, a key feature of our method is optimized cell yield and ability to expand edited cells for at least 3 weeks, enabling study of the progressive consequences of gene ablation. After optimization, we achieved 40–60% CRISPR-mediated gene knock-in and 100–200-fold nTreg expansion within 12 days. Previous studies using ssDNA, dsDNA, or AAV6 to knock-in genes in human Tregs reported variable HDR rates (10–50%) in cells cultured for 5–7 d, with limited reports of Treg yield or purity (Roth et al., 2018; Goodwin et al., 2020; Nguyen et al., 2020). Notably, we identified Treg-specific parameters critical for efficient CRISPR-mediated HDR editing. Dynabead-based pre-activation, commonly used for CRIPSR editing of total T cells (Eyquem et al., 2017; Liu et al., 2017; Ren et al., 2017; Chen et al., 2018; Roth et al., 2018; Nguyen et al., 2020), was not feasible for small numbers of Tregs (i.e., 1–4×10^5^ cells) because requisite bead removal prior to electroporation resulted in significant cell loss. We also found that inclusion of CD2 costimulation, either by anti-CD2 on tetramers or CD58 expression on aAPCs, resulted in superior Treg recovery after electroporation. This finding is consistent with previous studies which found that CD2 engagement on Tregs promotes cell survival (Kashiwakura et al., 2013). Finally, aAPC-based pre-activation substantially increased Treg expansion compared to cell-free activation reagents, likely by providing a more physiological form of stimulation (Cheung et al., 2018).

The incorporation of HDR-mediated gene knock-in with CRISPR editing has several advantages for functional studies of human Tregs. The majority of loss-of-function studies using CRISPR rely on error-prone DNA repair, which results in heterogeneous editing outcomes and on a single-cell basis can cause a range of functional phenotypes. Recent studies have uncovered compensatory pathways that can circumvent the intended CRISPR-mediated gene KO (El-Brolosy et al., 2019; Ma et al., 2019; Smits et al., 2019; Tuladhar et al., 2019). Moreover, wildtype cells may exert dominant and/or cell-extrinsic activity within a mixed pool of cells and obscure altered phenotypes; indeed, mixed bone-marrow chimeras of wildtype and *Foxp3*-deficient cells do not result in pathology (Fontenot, Gavin and Rudensky, 2003). Using HDR to introduce a reporter gene enables purification of cells that underwent precise editing and maximizes population homogeneity.

As FOXP3 is an X-linked gene subject to random X-inactivation (Tommasini et al., 2002; Carrel and Willard, 2005; Cotton et al., 2011), FOXP3 KO only required monoallelic reporter knock-in. If biallelic gene knock-in is required, then intensity of reporter gene expression (Dever et al., 2016; Oceguera-Yanez et al., 2016) and/or a double-reporter approach can be used (Roth et al., 2018). If using gene knock-in to achieve uniform gene KO, however, monoallelic reporter integration may be sufficient since cells which underwent HDR at one allele frequently exhibit error-prone DNA repair at the other allele (Zhang et al., 2014; Paquet et al., 2016; Wassef et al., 2017; Spiegel et al., 2019).

Most of our understanding of the role of Foxp3 in Treg biology comes from studies in which the gene is ablated prior to Treg lineage commitment. Thus far, three studies have investigated the effects of Foxp3 ablation in mature mouse or human Tregs. Using a self-inactivating Cre-expressing retroviral vector to achieve recombination at *Foxp3*, mouse Tregs lost suppressive capacity in a T cell adoptive transfer model, upregulated inflammatory cytokines, and exhibited an aberrant Treg phenotype at the transcript (*Ctla4, Entpd1* (CD39)) and protein levels (CD25) (Williams and Rudensky, 2007). Another group used lentiviral siRNA to achieve *FOXP3* knockdown in human Tregs, finding reduced in vitro suppressive function (Amendola et al., 2009). More recently, CRISPR-based FOXP3 KO in human Tregs revealed increased expression of IFN-γ, IL-4, and IL-2 and a trend towards reduced in vivo suppressive function (Schumann et al., 2020). In our system, FOXP3^HDR-KO^ Tregs did not significantly upregulate IL-2. This discrepancy may result from the timing of cytokine evaluation (5 days vs. 17 days post-editing in our study), which could affect Treg activation states and/or enable counter-regulatory mechanisms. In addition, these previous studies used total Tregs not separated into naive and memory subsets, which have known functional heterogeneity (Miyara et al., 2009).

Upon FOXP3 ablation, Tregs displayed subset-specific outcomes. FOXP3^HDR-KO^ mTregs exhibited greater dysfunction than their nTreg counterparts, upregulating cytokine expression to a greater extent and showing more pronounced loss of suppressive function. mTregs are a heterogeneous population and can acquire transcriptional programs mirroring that of Th cells (Duhen et al., 2012; Halim et al., 2017) which may become overactive in the absence of FOXP3 regulation. Indeed, the largest fraction of human mTregs are Th17-like Tregs co-expressing CCR6 and RORγt (Duhen et al., 2012; Halim et al., 2017; Boardman et al., 2020), consistent with the observation that FOXP3^HDR-KO^ mTregs strongly upregulated IL-17A secretion. It is unlikely that the exaggerated phenotype in FOXP3^HDR-KO^ mTregs is due to contamination of activated non-Tregs (Miyara et al., 2009), as control Cas9 mTregs maintained FOXP3 expression even after 2–3 weeks of culture.

FOXP3^HDR-KO^ nTregs exhibited a Th2-dominant phenotype: they upregulated secretion of IL-4, IL-5, and IL-13 and lost promoter DNA methylation at *IL5* and *IL13*, implying reprogramming at the epigenetic level. Although *IL5* and *IL13* DMRs did not overlap a FOXP3-bound region, they are regulated by GATA3, which is critical for both Treg homeostasis and Th2 differentiation and maintenance (Wang, Su and Wan, 2011; Wohlfert et al., 2011; Rudra et al., 2012). Tregs from mice with attenuated Foxp3 expression, or from mice with a loss-of-function FOXP3 mutation (M370I) found in an IPEX patient, acquire a GATA3-dependent Th2 effector program (Wan and Flavell, 2007; Van Gool et al., 2019). Given that GATA3 binds DNA in a methylation-insensitive manner and remodels chromatin at Th2 cytokine genes (Yamashita et al., 2004; Yin et al., 2017), it is tempting to speculate that, in nTregs, a major role of FOXP3 is to actively prevents these processes.

Functionally, FOXP3^HDR-KO^ Tregs progressively lost suppressive capacity in vitro, particularly towards CD8^+^ T cells, suggesting that suppression of CD8^+^ rather than CD4^+^ T cell proliferation may be a more sensitive measurement of changes in Treg function. What underlies this loss of suppressive function? Decades of work suggest that Tregs likely use multiple mechanisms to suppress T cell activity in a cell contact- or proximity-dependent fashion (Schmidt, Oberle and Krammer, 2012), yet expression of proteins involved in the Treg repertoire of suppressive mechanisms, including CD25, CTLA-4, CD39, and LAP, remained unchanged on FOXP3-ablated Tregs. Whether Tregs utilize distinct pathways to suppress CD4^+^ T cell vs. CD8^+^ T cell proliferation requires further investigation.

Despite their dysfunction, FOXP3^HDR-KO^ Tregs retained a Treg protein phenotype and DNA methylation signature. In mouse Treg development, precursor cells acquire Treg-type DNA demethylation at several regions independently of Foxp3, and Foxp3 exploits a pre-existing enhancer landscape (Ohkura et al., 2012; Samstein et al., 2012). In addition, FOXP3-binding regions in mice and humans do not substantially overlap Treg-specific DMRs (Morikawa et al., 2014; Ohkura et al., 2020). In our system, only a fraction of Treg-specific DMRs were affected by FOXP3 ablation, and the majority of FOXP3 KO-induced DMRs did not overlap FOXP3-binding regions identified by ChIP-chip (Sadlon et al., 2010). Overall, our findings suggest that, beyond Treg development, FOXP3 has a limited role in the maintenance of established human Treg identity. This interpretation is consistent with a recent study showing that Foxp3 affects mouse Treg chromatin accessibility indirectly via other transcription factors (van der Veeken et al., 2020).

Amongst the limited changes in methylation, FOXP3 KO predominantly induced DNA demethylation with an enrichment in regions encoding bZIP (AP-1) binding motifs. CpG methylation is heritably maintained by the DNA methyltransferase DNMT1, which can exert region-specific effects when bound to a transcription factor (Hervouet et al., 2018). Accordingly, Foxp3 complexes with DNMT1 (Wang et al., 2013), though the functional consequences of this interaction remain undefined. A recent study reported that mice with a Treg-specific inducible deletion of the DNMT1 adaptor *Uhrf1* resulted in selective CpG demethylation and destabilization of mature mouse Tregs (Helmin et al., 2020). Analogously, we found that FOXP3-induced Treg dysfunction was associated with DNA demethylation in specific regions which were enriched for AP-1 binding sites. AP-1 family members are thought to prepare sites for Foxp3 binding in Treg precursors (Samstein et al., 2012) and control the effector and tissue-specific Treg programs (DiSpirito et al., 2018; Mijnheer et al., 2020). On the other hand, Foxp3 opposes AP-1 activity by competing for binding with NFAT (Wu et al., 2006; Lee, Gao and Fang, 2008). Collectively, these results suggest that FOXP3 KO-induced DNA demethylation may enable aberrant AP-1 activity.

The ability to flexibly knock-out and knock-in genes in human Tregs in a way that enables purification of successfully targeted cells and expansion for mechanistic studies and/or therapy opens many new possibilities. The finding that deletion of the Treg master transcription factor did not substantially alter the characteristic Treg phenotype or epigenome, and that significant functional effects were only evident after prolonged culture, underscores the need for further investigation into how the Treg lineage is maintained in humans. Overall, optimized CRISPR-mediated gene knock-in will enable the design and generation of Tregs with tailored therapeutic effects.

## Acknowledgements

We thank Rosa Bacchetta’s lab (Stanford University) for helpful discussions regarding AAV6 purification and the National Research Council for providing AAV6-CMV-GFP. We thank Julie MacIsaac and Oscar Urtatiz (University of British Columbia; UBC) for performing DNA methylation array assays. We thank UBC Nucleic Acid Protein Service and the BC Children’s Hospital Research Institute (BCCHR) DNA Sequencing and Flow Core Facilities for technical support. This work was supported by grants from the Canadian Institutes of Health Research (CIHR) (FDN-154304 to MKL). AJL is supported by a CIHR Doctoral Research Award and MKL receives a BCCHR salary award.

## Author Contributions

Conceptualization: AJL, MKL. Methodology: AJL, DTSL, JG, AMP. Investigation: AJL, DTSL, JG, PU. Formal analysis: AJL, DTSL. Writing—Original Draft: AJL. Writing—Review & Editing: AJL, DSTL, JG, AMP, MSK, MKL. Supervision: MSK, MKL. Funding Acquisition: MKL

## Methods

### CRISPR and HDR template design and assembly

*CD226*- and *FOXP3*-targeting gRNAs were synthesized as chemically modified crRNAs (IDT) and duplexed at a 1:1 molar ratio with tracrRNA (IDT) by denaturation for 5 min at 95°C followed by gradual cooling to room temperature. The resulting gRNA was complexed with Cas9 (QB3 Macrolab) at a 2:1 molar ratio for 10 min at room temperature. gRNA target sequences: CD226 (5′-GTTAAGAGGTCGATCTGACG-3′), FOXP3 CR1 (5′-AGGACCCGATGCCCAACCCC-3′), FOXP3 CR2 (5′-GGGCCGAGATCTTCGAGGCG-3′), FOXP3 CR3 (5′-GCAGCTGCGATGGTGGCATG-3′), FOXP3 CR4 (5′-TGCCCCCCAGCTCTCAACGG-3′), FOXP3 CR5 (5′-CCCACCCACAGGGATCAACG-3′). HDR designs contained homology arms of at least 800 bp flanking a promoterless or EF-1α-driven transgene marker (sfGFP or ΔNGFR). Fragments were PCR-amplified from genomic DNA, plasmids, or synthesized (IDT gBlocks or GenScript GenParts), then cloned by Gibson assembly into an AAV transfer plasmid between two ITRs.

### AAV6 production, purification, and titration

For recombinant AAV6 production, HEK-293T/17 cells (ATCC CRL-11268) were co-transfected by calcium phosphate with an adenoviral pHelper plasmid, a pAAV6-Rep-Cap plasmid (both Cell Biolabs), and an ITR-flanked transfer plasmid at a 1:1:1 molar ratio. Cells were harvested after 3 d and AAV6 was purified with a filtration-based kit (Takara AAVpro Purification Kit) per the manufacturer’s protocol. AAV6 vector genomes were titrated by ITR-specific quantitative PCR (Takara AAVpro Titration Kit Ver.2) per the manufacturer’s protocol.

### Cell isolation and culture

Human blood sample collection from healthy adults was performed in accordance with protocols approved by the University of British Columbia Clinical Research Ethics Board and Canadian Blood Services. CD3^+^ T cells or CD4^+^ T cells were isolated via Lymphoprep and RosetteSep (both STEMCELL Technologies). For Treg isolation, CD4^+^ T cells were enriched with CD25 MicroBeads II (Miltenyi Biotec) before flow sorting on a MoFlow Astrios (Beckman Coulter) or FACSAria IIu (BD Biosciences). Sorting strategies: total Tregs (CD4^+^CD25^hi^CD127^lo^), naive Tregs (nTreg; CD4^+^CD25^hi^CD45RA^+^CD127^lo^), memory Tregs (mTreg; CD4^+^CD25^hi^CD45RA^−^CD127^lo^), and conventional T cells (Tconv; CD4^+^CD25^lo^CD127^hi^). Unless otherwise indicated, all cells were cultured at 37°C, 5% CO2 in X-VIVO 15 (Lonza) supplemented with 5% (v/v) human serum (WISENT), 1% (v/v) penicillin-streptomycin (Gibco), 2 mM GlutaMAX (Gibco), and 15.97 mg/L phenol red (Sigma-Aldrich). During optimization, in some cases as indicated, CTS OpTmizer T Cell Expansion SFM (Gibco) was used in place of X-VIVO 15, human serum, and phenol red.

### Human Treg expansion with CRISPR/HDR-mediated gene knock-in

In all cases during pre-activation and expansion, media was supplemented with IL-2 (Proleukin; 1000 IU/ml for Tregs, 100 IU/ml for Tconvs or CD4^+^ T cells). Tregs, Treg subsets, Tconvs, and CD4^+^ T cells were pre-activated with either CD3/CD28/CD2 tetrameric antibody complexes (ImmunoCult by STEMCELL Technologies) or 1:1 aAPCs for 5 d. aAPCs were L cells (ATCC CRL-2648) expressing CD32, CD58, and CD80, gamma-irradiated (75 Gy), and loaded with and anti-CD3 (OKT3, 0.1 μg/ml; University of British Columbia Antibody Lab) (de Waal Malefyt et al., 1993). Media and IL-2 were replenished every 2–3 d.

Cells were electroporated with the Neon Transfection 10 μL Kit (Invitrogen) per the manufacturer’s protocol. Briefly, cells were washed twice with PBS, resuspended in Buffer T (≤20×10^6^ cells/mL) with Cas9 or RNP (40 pmol gRNA + 20 pmol Cas9 per transfection), electroporated at 1400 V / 30 ms / 1 pulse, then immediately transferred into prewarmed antibiotic-free media containing AAV6 at the specified vg/cell and simultaneously expanded with aAPCs and IL-2 for 7 d as above. After 7 d expansion (total 12 d), cells were flow-sorted as sfGFP^+^, then rested overnight in reduced IL-2 (100 IU/ml for Tregs, none for Tconvs) before functional assays; in some cases, flow-sorted cells were further expanded for 7 d with aAPCs and IL-2 as above before use in assays. Cell counts at 5 d were determined by trypan blue (Gibco); cell counts at 12 d and 19 d were determined by ViaStain Acridine Orange/Propdium Iodide Staining Solution (Nexcelom).

For experiments investigating the role of FOXP3 in Treg biology, we pre-activated cells with CD3/CD28/CD2 tetramers followed by aAPC-based expansions, for two reasons. First, Treg yield by day 19 of expansion was comparable to a fully aAPC-based expansion (**Figure 1D**). Second, we found that repetitive (3×) aAPC-based stimulation occasionally resulted in spontaneous FOXP3 loss (data not shown), which could be circumvented by using a weaker reagent to pre-activate Tregs, i.e., CD3/CD28/CD2 tetramers.

During optimization, cells were alternatively pre-activated with IL-2 as above and one of: 1:1 anti-CD3/anti-CD28-coated beads (Gibco Dynabeads Human T-Expander or Human Treg Expander), CD3/CD28 tetrameric antibody complexes (ImmunoCult by STEMCELL Technologies), or a combination of plate-bound anti-CD3 (OKT3, 10 μg/ml in 0.1 M Trizma HCl; Sigma-Aldrich) with soluble anti-CD28 (CD28.2, 4 μg/ml; BD Biosciences). Dynabead-activated cells were magnetically de-beaded immediately prior to electroporation. Divergent T cell pre-activation conditions and Neon electroporation parameters are specified as applicable. In some cases, HDR Enhancer (IDT, v1) was added to AAV6-containing media prior to electroporation at the indicated concentrations.

### Evaluation of genome editing

To assess indel formation after CRISPR editing, genomic DNA was extracted from cells 3 d after electroporation using a QIAamp DNA Mini Kit or Micro Kit (QIAGEN) per the manufacturer’s protocols. The region flanking the gRNA cut site was PCR-amplified, Sanger-sequenced, and chromatograms uploaded to the ICE analysis tool (Synthego; https://ice.synthego.com/) (Hsiau et al., 2019). Primer sequences for FOXP3 CR1 and FOXP3 CR3: FWD (5′-CTAGAGCTGGGGTGCAACTATG-3′), REV* (5′-TCTTCTCTTGTCACATGGGGATG-3′). Primer sequences for FOXP3 CR5: FWD* (5′-GAATGGCCGTCTTTAAGCTTCTC-3′), REV (5′-TTATTGGGATGAAGCCTGAGCTG-3′). An asterisk (*) denotes the primer used for Sanger sequencing.

### T cell suppression assay

Expanded Tregs and ex vivo-isolated allogeneic CD3^+^ T cells (responder cells) were labelled with Cell Proliferation Dye (Invitrogen) eFluor 670 and eFluor 450, respectively, then cocultured at the indicated ratios and activated with anti-CD3/anti-CD28-coated beads (Gibco Dynabeads T-Expander; 1:16 bead:responder cells) for 4 d. Proliferation of CD4^+^ and CD8^+^ cells within the responder-cell fraction was determined by dilution of Cell Proliferation Dye; responder cells activated with beads served as positive controls. Percent suppression of proliferation was calculated as: (1 − (division index of sample / division index of positive control)) * 100.

### Assessment of cytokine production

Expanded Tregs or Tconvs were activated with IL-2 (100 IU/mL) and 1:1 anti-CD3/anti-CD28-coated beads (Gibco Dynabeads T-Expander) for 4 d. For secreted cytokines, supernatants were collected, cytokines measured by cytometric bead array (BioLegend LEGENDplex Human Th Panel 13-plex), and data analysed with Qognit software (BioLegend) per the manufacturer’s protocols. For intracellular cytokine expression, cells were restimulated with PMA (10 ng/ml), ionomycin (500 μg/ml), and brefeldin A (10 μg/ml; all Sigma-Aldrich) for an additional 4 h.

### Flow cytometry

Antibodies are listed in **Table S2**. Cells were stained for surface proteins in PBS (Gibco) or Brilliant Stain Buffer (BD Biosciences) for 20 min at room temperature; Fixable Viability Dye (Invitrogen) or 7-AAD (BioLegend) was used to exclude dead cells. For detection of intracellular proteins, cells were fixed and permeabilized with the eBioscience Foxp3 / Transcription Factor Staining Buffer Set (Invitrogen) for 40 min at room temperature, then stained for intracellular proteins for 40 min at room temperature. As applicable, Brilliant Stain Buffer Plus (BD Biosciences) was included during intracellular staining. Samples were acquired on an LSRFortessa X-20, FACSymphony A5 (both BD Biosciences), or Cytoflex (Beckman Coulter), and data were analysed with FlowJo software (BD Biosciences; v10.7). All live single cells were first gated as CD4^+^, except for the T cell suppression assay in which live single cells were gated as CD4^+^ or CD8^+^. Percent protein KO was calculated as: (1 – (% marker^+^ of RNP-edited sample / % marker^+^ of negative control)) * 100.

### Genome-wide DNA methylation profiling

FOXP3^HDR-KO^ nTregs, Cas9 nTregs, and Tconvs expanded for 20 d from 5 individuals were used for genome-wide DNA methylation profiling. Genomic DNA was isolated with an AllPrep DNA/RNA Mini Kit (QIAGEN), bisulfite-converted with an EZ DNA Methylation Kit (Zymo Research), hybridized to Infinium MethylationEPIC BeadChips (Illumina), and imaged using an iScan System (Illumina) per the manufacturers’ protocols. Raw red and green intensity data (IDAT) were imported into the R statistical environment (v4.0.2) with RStudio (v1.3.1093), converted into methylated and unmethylated signals, normalized using noob (Triche et al., 2013) and functional normalization (Fortin et al., 2014) by preprocessFunnorm in minfi (v1.34.0) and by BMIQ (Teschendorff et al., 2013) in wateRmelon (v1.32.0), and corrected for batch effects (BeadChip ID and position) using ComBat in sva (v3.36.0). Probes with a detection p-value > 0.01 or a bead count < 3 in at least one sample were discarded. Probes previously characterized to have low-quality genomic mapping, cross-reactive or polymorphic sequences, or type I probes targeting a common SNP that could cause a colour channel switch were discarded (Zhou, Laird and Shen, 2017) (https://zwdzwd.github.io/InfiniumAnnotation). Because the cohort contained cells from males and females, probes targeting X/Y chromosomes were discarded. The total remaining CpG sites after preprocessing was 702,481. Results were visualized as beta values, which range from 0 (unmethylated) to 1 (fully methylated).

Principal component analysis was performed using prcomp. Differential methylation analysis for all paired comparisons was performed with logit-transformed beta values (M values), considered more statistically robust (Du et al., 2010), using limma (v3.44.3). CpG sites with a mean methylation (beta) difference > 0.05 and Benjamini-Hochberg-corrected p-value < 0.05 were considered differentially methylated. DMRs were then identified by DMRcate (v2.2.3) (Peters et al., 2015) using a bandwidth of 1000 nt (lambda = 1000) and a scaling factor of 2 (C = 2). Regions with contiguous, differentially methylated CpG sites within lambda nucleotides, a region mean methylation (beta) difference > 0.05, and Fisher’s multiple-comparison statistic < 0.05 were considered significantly differentially methylated.

To identify differentially methylated regions that overlapped a FOXP3-binding region, a publicly available human FOXP3 whole-genome ChIP-chip tiled array dataset was used (Sadlon et al., 2010). The genomic coordinates of FOXP3-binding sites identified by model-based analysis of tiling-arrays were lifted over from hg18 to hg19, then compared to the DMRs identified above. A DMR was considered overlapping a FOXP3-binding region if there was an overlap of at least 1 nt.

HOMER (Heinz et al., 2010) was used for transcription factor binding motif enrichment analysis. The coordinates of demethylated DMRs were scanned using findMotifsGenome.pl with default parameters, exact (given) fragment size, and “-mask” to mask repeat sequences. Background sequences with matching GC content were randomly selected as controls. Demethylated DMRs were searched for overrepresented motifs with lengths of 8, 10, and 12 bp relative to the background sequences. De novo motifs with p-value < 1×10^−12^ were considered enriched.

### Statistical analysis

Normality was assumed. Statistical significance between more than two groups was determined by matched 1-way ANOVA or matched 2-way mixed-effects model and Dunnett’s or Tukey’s multiple comparisons tests, as appropriate. Significance between two groups was determined by paired t-test; for lognormal distributions, ratio paired t-test or Mann-Whitney test was used, as appropriate. For AAV6 dose response experiments, areas under the curve were used to determine significance. For suppression of proliferation, areas under the curve were used to determine significance, taking into account all degrees of freedom, as previously described (Akimova et al., 2016). The n values used to calculate statistics are defined in the figure legends; significance (p < 0.05 was considered significant) is indicated within the figures. Analysis was performed using Prism software (GraphPad; v9.0.0). Statistical testing of DNA methylation patterns was performed in R, as described in the preceding section.

