## Supplemental Figures & Tables for "Optimized CRISPR-mediated gene knock-in reveals FOXP3-independent control of human Treg identity"

### Supplemental Information

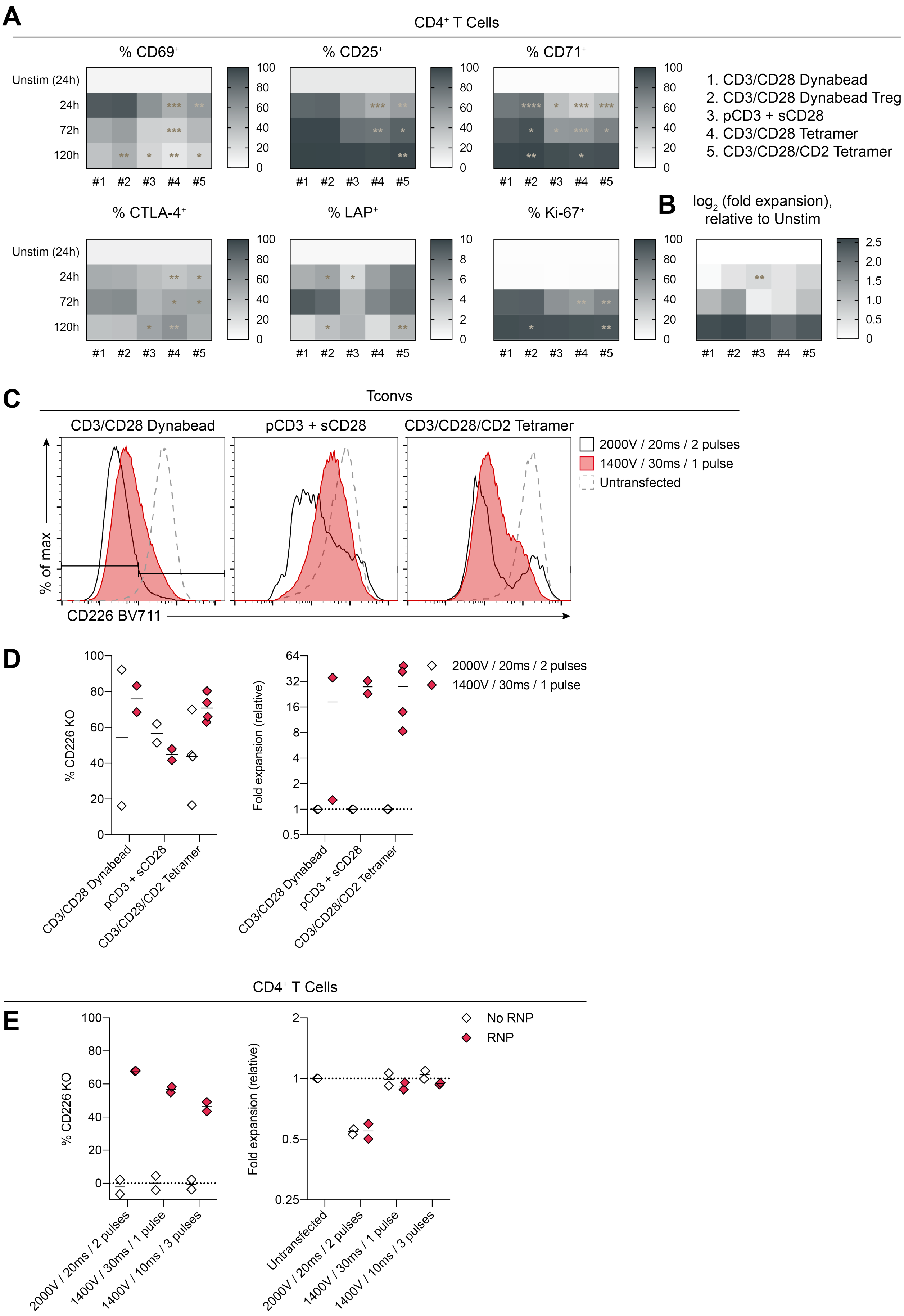

***Figure S1. Optimization of T cell pre-activation and CRISPR electroporation, related to Figure 1.***

(**A-B**) $CD4^+$ T cells were activated with the indicated reagent and analyzed at the indicated time for (**A**) expression of CD69, CD25, CD71, CTLA-4, LAP, and Ki-67 and (**B**) fold expansion relative to 24 h-unstimulated cells (n=6, 2 experiments). (**C-D**) Conventional T cells (Tconv; $CD4^+CD25^{lo}CD127^{hi}$) were pre-activated with the indicated reagent (5 d), electroporated with a *CD226*-targeting gRNA, and expanded with artificial APCs (aAPCs) (3 d) (n=2–4, 2 experiments). (**C**) Representative CD226 expression. (**D**) Left: percent CD226 KO, relative to untransfected cells. Right: fold expansion, relative to 2200 V / 20 ms / 2 pulses within each activation agent. (**E**) $CD4^+$ T cells were pre-activated with CD3/CD28 tetramers (25% dose) in CTS OpTmizer (3 d), electroporated with or without a *CD226*-targeting gRNA, and expanded with aAPCs (3 d) (n=2, 1 experiment). Left: Percent CD226 KO, relative to untransfected cells. Right: Fold expansion, relative to untransfected cells. (**A-B**) depicts mean; solid lines and dots in (**D-E**) represent means and individual donors, respectively. Each dot in (**E**) depicts the average of 2 technical replicates. Significance in (**A-B**) determined by matched 2-way mixed-effects model with Geisser-Greenhouse correction and Dunnett's multiple comparisons test, with all comparisons to #1 CD3/CD28 Dynabeads.

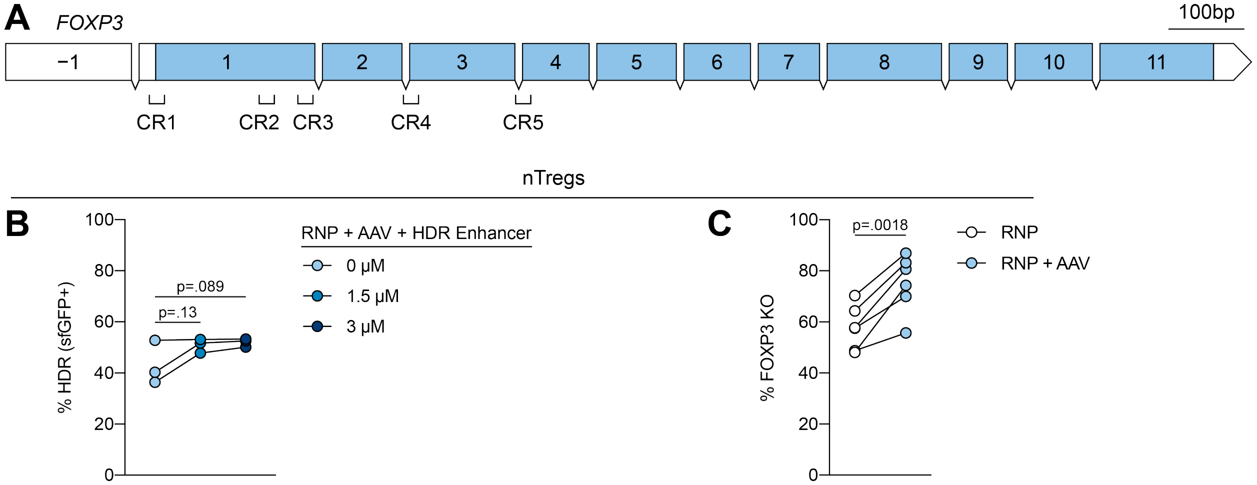


***Figure S2. Strategies to increase CRISPR and HDR efficiency in human Tregs, related to Figure 2.***

(**A**) Schematic diagram of human *FOXP3* gene with exon numbers and gRNA locations. Introns not to scale. (**B**) Naive Tregs (nTreg; CD4$^{+}$CD25$^{hi}$CD127$^{lo}$CD45RA$^{+}$) were pre-activated with CD3/CD28/CD2 tetramers (5 d), electroporated with FOXP3 CR3 gRNA, then simultaneously transduced with AAV6-HDR and treated with HDR Enhancer (IDT, v1) at the indicated concentrations, and expanded with aAPCs (6 d). sfGFP expression (HDR) (n=3, 2 experiments). (**C**) nTregs were pre-activated and electroporated as in (**B**), then transduced with AAV6-HDR (or not) and expanded with aAPCs (6 d). (**C**) Percent FOXP3 KO, relative to Cas9 cells, by flow cytometry (n=6, 3 experiments). (**B-C**) depict individual donors. Significance determined by matched 1-way ANOVA with Dunnett's multiple comparisons test for (**B**) and paired t-test for (**C**).

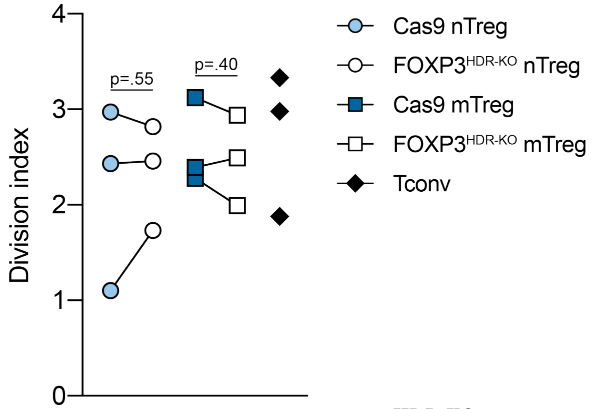


***Figure S3. Cas9 and FOXP3[HDR-KO] Tregs proliferate similarly upon strong TCR stimulation, related to Figure 4.***

Cas9 and FOXP3[HDR-KO] nTregs and mTregs were expanded for 13 d total, then TCR-activated (4 d) (n=3, 3 experiments). Division index depicted. Dots represent individual donors. Significance determined by paired t-test; nTregs and mTregs evaluated separately. Tconv for reference only.

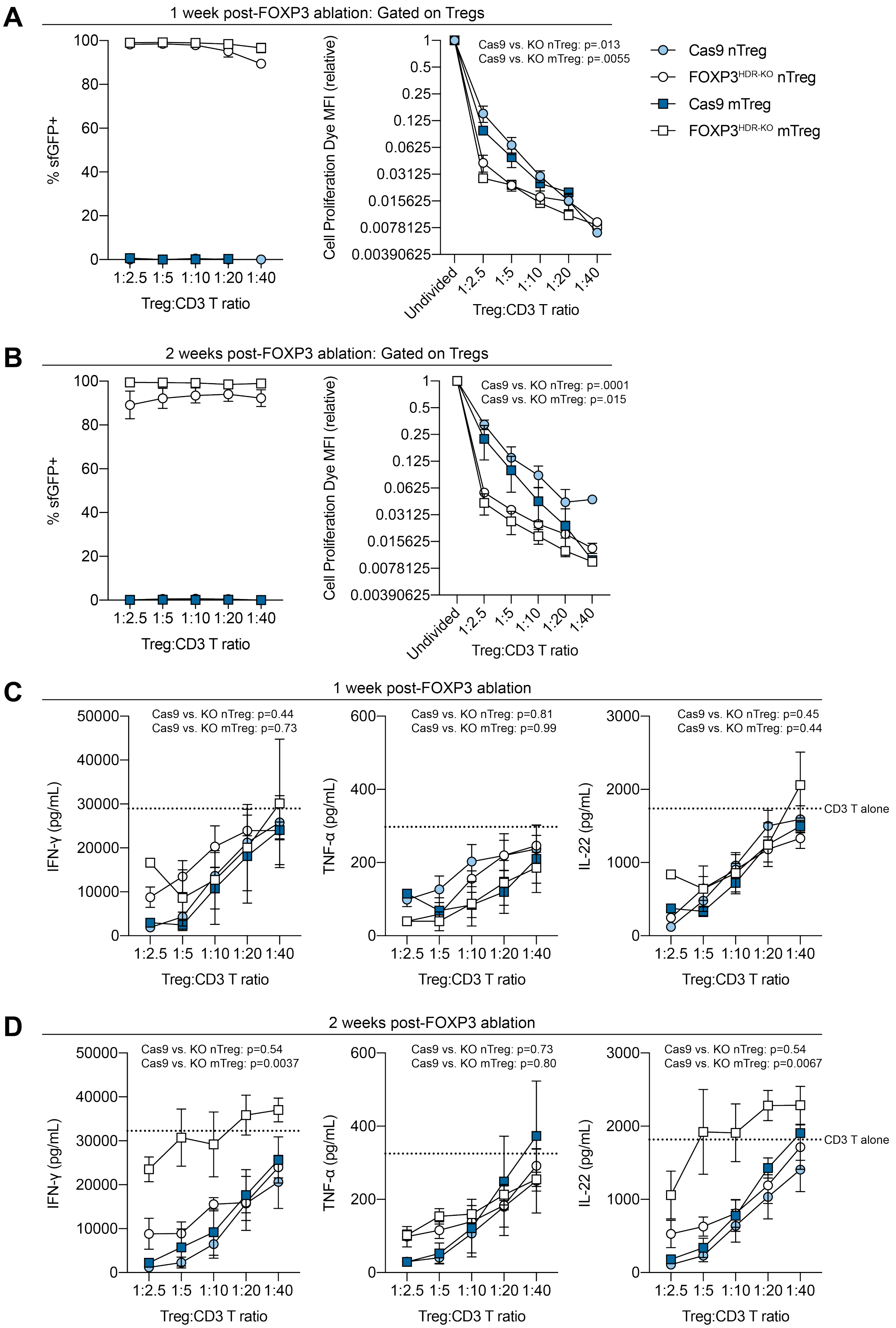
A
1 week post-FOXP3 ablation: Gated on Tregs
% sfGFP+
Treg:CD3 T ratio
Cell Proliferation Dye MFI (relative)
Cas9 vs. KO nTreg: p=.013
Cas9 vs. KO mTreg: p=.0055
Undivided
Cas9 nTreg
FOXP3HDR-KO nTreg
Cas9 mTreg
FOXP3HDR-KO mTreg
B
2 weeks post-FOXP3 ablation: Gated on Tregs
Cas9 vs. KO nTreg: p=.0001
Cas9 vs. KO mTreg: p=.015
C
1 week post-FOXP3 ablation
IFN-γ (pg/mL)
TNF-α (pg/mL)
IL-22 (pg/mL)
Cas9 vs. KO nTreg: p=0.44
Cas9 vs. KO mTreg: p=0.73
Cas9 vs. KO nTreg: p=0.81
Cas9 vs. KO mTreg: p=0.99
Cas9 vs. KO nTreg: p=0.45
Cas9 vs. KO mTreg: p=0.44
CD3 T alone
D
2 weeks post-FOXP3 ablation
Cas9 vs. KO nTreg: p=0.54
Cas9 vs. KO mTreg: p=0.0037
Cas9 vs. KO nTreg: p=0.73
Cas9 vs. KO mTreg: p=0.80
Cas9 vs. KO nTreg: p=0.54
Cas9 vs. KO mTreg: p=0.0067
CD3 T alone

***Figure S4. FOXP3$^{HDR\text{-}KO}$ Tregs are more proliferative and are less able to suppress cytokine production in a T cell suppression assay, related to Figure 5.***

Cas9 and FOXP3$^{HDR\text{-}KO}$ nTregs and mTregs were expanded for (**A, C**) 13 d or (**B, D**) 20 d total (n=2–4 nTreg, n=1–3 mTreg, 3 experiments). Cells were cocultured with allogeneic $CD3^+$ T cells (responder cells) at the indicated ratios and activated with CD3/CD28 Dynabeads (4 d). (**A-B**) Tregs were gated by differential staining of Cell Proliferation Dye. Left: sfGFP expression. Right: Treg proliferation, by MFI of Cell Proliferation Dye relative to the MFI of that of the undivided fraction. (**C-D**) Supernatant concentrations of IFN-γ, TNF-α, and IL-22; dotted line represents cytokine concentrations from Dynabead-activated $CD3^+$ T cells cultured alone. (**A-B**) depicts mean±SEM. Significance in (**A-B**, right and **C-D**) determined by t-test of the areas under the curve; nTregs and mTregs evaluated separately. MFI, geometric mean fluorescence intensity.

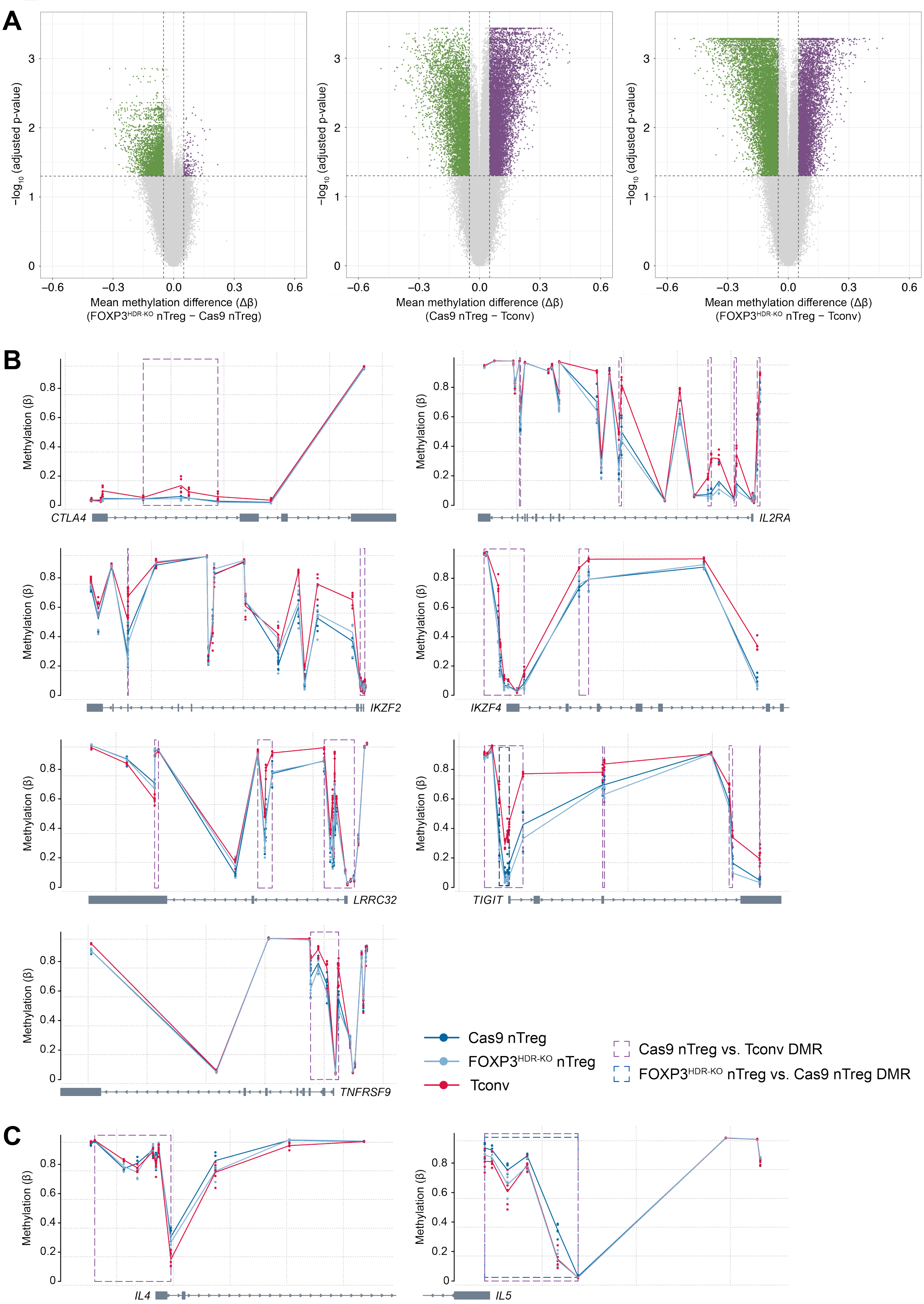
A
−log10 (adjusted p-value)
Mean methylation difference (Δβ)
(FOXP3HDR-KO nTreg − Cas9 nTreg)
Mean methylation difference (Δβ)
(Cas9 nTreg − Tconv)
Mean methylation difference (Δβ)
(FOXP3HDR-KO nTreg − Tconv)
B
Methylation (β)
CTLA4
IL2RA
IKZF2
IKZF4
LRRC32
TIGIT
TNFRSF9
Cas9 nTreg
FOXP3HDR-KO nTreg
Tconv
Cas9 nTreg vs. Tconv DMR
FOXP3HDR-KO nTreg vs. Cas9 nTreg DMR
C
IL4
IL5

***Figure S5. FOXP3-dependent and -independent changes in nTreg DNA methylation, related to Figure 7.***

(**A-C**) Genome-wide DNA methylation was assessed in 20 d-expanded FOXP3$^{HDR\text{-}KO}$ nTregs, Cas9 nTregs, and Tconvs (n=5). (**A**) Volcano plot of differentially methylated CpG sites for FOXP3$^{HDR\text{-}KO}$ nTregs vs. Cas9 nTregs (left), Cas9 nTregs vs. Tconvs (middle), and FOXP3$^{HDR\text{-}KO}$ nTregs vs. Tconvs (right). Differentially methylated CpG sites are defined as having a mean methylation (beta) difference > 0.05 and Benjamini-Hochberg-corrected p-value < 0.05. Gene-associated DMRs for (**B**) *CTLA4*, *IL2RA* (CD25), *IKZF2* (Helios), *IKZF4* (Eos), *LRRC32* (GARP), *TIGIT*, *TNFRSF9* (CD137), (**C**) *IL4*, and *IL5*. DMRs are defined as having a region mean methylation (beta) difference > 0.05 and Fisher's multiple-comparison statistic < 0.05. Dots in (**A**) represent CpG sites, with differentially methylated sites delineated by the dotted lines. Dots in (**B-C**) represent individual samples at each CpG site. DMRs in (**B-C**) are delineated by dotted boxes (purple: Cas9 nTreg vs. Tconv DMR; dark blue: FOXP3$^{HDR\text{-}KO}$ nTreg vs. Cas9 nTreg DMR).

***Table S1. Differentially methylated CpG sites, regions, and overlaps with FOXP3-binding regions, related to Figure 7.***

CpG sites with a mean methylation (beta) difference > 0.05 and Benjamini-Hochberg-adjusted p-value < 0.05 were considered differentially methylated (DM). Regions identified by DMRcate were considered differentially methylated (DMR) if the region mean methylation (beta) difference > 0.05 and Fisher's multiple-comparison statistic < 0.05. FOXP3-binding regions inferred from a publicly available human FOXP3 whole-genome ChIP-chip dataset (Sadlon et al., 2010).

| | $FOXP3^{HDR-KO}$ nTreg vs. Cas9 nTreg | Cas9 nTreg vs. Tconv | $FOXP3^{HDR-KO}$ nTreg vs. Tconv |
|---|---|---|---|
| Differentially methylated CpG sites (#) | 5151 | 18 552 | 18 961 |
| More-methylated CpG sites (# (% of DM sites)) | 168 (3.26%) | 11 894 (64.1%) | 7466 (39.4%) |
| Less-methylated CpG sites (# (% of DM sites)) | 4983 (96.7%) | 6658 (35.9%) | 11 495 (60.6%) |
| Differentially methylated regions (#) | 471 | 2326 | 2633 |
| More-rmethylated regions (# (% of DMRs)) | 14 (2.97%) | 1401 (60.2%) | 962 (36.5%) |
| Less-methylated regions (# (% of DMRs) | 457 (97.0%) | 925 (39.8%) | 1671 (63.5%) |
| Differentially methylated regions overlapping a FOXP3-binding region (# (% of DMRs)) | 99 (21.0%) | 422 (18.1%) | 467 (17.7%) |

***Table S2. List of antibodies used for flow cytometry, related to Methods.***

| Target | Clone | Fluorophore | Company |
|---|---|---|---|
| CD3 | UCHT1 | BB515 | BD Biosciences |
| CD4 | OKT4 | BV421 | BioLegend |
| CD4 | RPA-T4 | BV605 | BD Biosciences |
| CD4 | RPA-T4 | PE | eBioscience |
| CD4 | RPA-T4 | V500 | BD Biosciences |
| CD4 | SK3 | BUV395 | BD Biosciences |
| CD4 | SK3 | BUV496 | BD Biosciences |
| CD8 | HIT8a | PE | BD Biosciences |
| CD25 | 2A3 | BUV395 | BD Biosciences |
| CD25 | 4E3 | PE | Miltenyi Biotec |
| CD25 | BC96 | BV421 | BioLegend |
| CD39 | A1 | BV421 | BioLegend |
| CD39 | eBioA1 | PerCP-eF710 | eBioscience |
| CD45RA | HI100 | FITC | eBioscience |
| CD69 | FN50 | BV711 | BioLegend |
| CD71 | M-A712 | BV786 | BD Biosciences |
| CD127 | eBioRDR5 | eF450 | eBioscience |
| CD152 (CTLA-4) | BNI3 | APC | BD Biosciences |
| CD152 (CTLA-4) | BNI3 | BV421 | BioLegend |
| CD152 (CTLA-4) | BNI3 | BV786 | BD Biosciences |
| CD184 (CXCR3) | G025H7 | BV421 | BioLegend |
| CD184 (CXCR3) | G025H7 | PE | BioLegend |
| CD194 (CCR4) | L291H4 | BV510 | BioLegend |
| CD196 (CCR6) | G034E3 | BV785 | BioLegend |
| CD226 | DX11 | BV711 | BD Biosciences |
| CD271 (NGFR) | C40-1457 | BV421 | BD Biosciences |
| CD271 (NGFR) | C40-1457 | PE-Cy7 | BD Biosciences |
| CD279 (PD-1) | eBioJ105 | PerCP-eF710 | eBioscience |
| CD279 (PD-1) | EH12.1 | BUV737 | BD Biosciences |
| CD279 (PD-1) | EH12.1 | PE-Cy7 | BD Biosciences |
| CD336 (Tim-3) | 344328 | BB700 | BD Biosciences |
| CD336 (Tim-3) | F38-2E2 | APC | BioLegend |
| FOXP3 | 236A/E7 | PE | eBioscience |
| FOXP3 | 236A/E7 | PE-Cy7 | eBioscience |
| Helios | 22F6 | AF647 | BioLegend |
| Helios | 22F6 | eF450 | eBioscience |
| HLA-DR | G46-6 | BV480 | BD Biosciences |
| HLA-DR | L243 | BV510 | BioLegend |
| IFN-γ | B27 | BUV395 | BD Biosciences |
| IL-2 | MQ1-17H12 | BUV737 | BD Biosciences |
| IL-17A | BL168 | BV711 | BioLegend |
| IL-17A | eBio64DEC17 | PE | eBioscience |
| Ki-67 | 20Raj1 | FITC | eBioscience |

| | | | |
|---|---|---|---|
| Ki-67 | B56 | BUV805 | BD Biosciences |
| Ki-67 | B56 | BV421 | BD Biosciences |
| LAP | FNLAP | PE | eBioscience |
| LAP | FNLAP | PE-Cy7 | eBioscience |
| TIGIT | A15153G | AF647 | BioLegend |
| TIGIT | MBSA43 | PE-Cy7 | eBioscience |
